# Is metabolism spatially optimized? Structural modeling of consecutive enzyme pairs reveals no evidence for spatial optimization of catalytic site proximity

**DOI:** 10.64898/2026.03.24.713955

**Authors:** Joaquin Algorta, Dirk Walther

## Abstract

Metabolic pathways are often hypothesized to benefit from the spatial organization of enzymes, facilitating substrate transfer through mechanisms such as metabolic channeling or metabolon formation. However, it remains unclear whether the spatial proximity of catalytic sites represents a general organizational principle of metabolism or is restricted to specific pathways. Here, we investigate whether consecutive enzymes in metabolic pathways, when physically interacting, exhibit structurally optimized arrangements that minimize distances between their catalytic sites, thereby increasing metabolite transfer efficiency from one enzyme to the next. We first evaluated the ability of current protein-protein interaction prediction methods, including AlphaFold2, AlphaFold3, ESMFold, and HDOCK, to model weak and transient interactions using a benchmark dataset of 112 low-affinity protein dimers from PDBbind. AlphaFold-based approaches performed best in recovering correct interaction geometries, while ESMFold showed limited performance. We further assessed several confidence metrics and identified ipTM, ipSAE, and VoroIF-GNN as the most informative predictors of correct interaction conformations. In addition to simple Euclidean distance metrics, we developed a computational procedure to estimate shortest accessible space paths between catalytic sites in predicted enzyme-enzyme complexes. Applying this framework to 107 consecutive enzyme pairs in *E.coli* revealed an increased tendency for consecutive enzymes to interact, but no systematic evidence that interacting enzymes position their catalytic sites in spatially optimized configurations. In the predicted complex conformations, catalytic sites tend not to be positioned closer than expected at random. The developed computational workflow provides a general framework for analyzing structural aspects of metabolic organization.

## Introduction

Cellular metabolism comprises a dense network of enzymatic reactions that has evolved under multiple, partly competing constraints, including catalytic efficiency, robustness, regulatory control, and resource allocation^1–4^. Beyond these factors, it has long been proposed that the spatial organization of metabolic enzymes may contribute to metabolic performance and kinetic optimization^5,6^. Such spatial effects are often discussed under concepts such as metabolic channeling, enzyme colocalization, or metabolon formation, and have been reported in selected pathways and organisms^7–9^. However, the extent to which these phenomena reflect general organizational principles of metabolism rather than pathway-specific adaptations remains unclear.

Early experimental evidence for metabolic channeling arose from kinetic measurements, isotope dilution experiments, and experimental detection of physical enzyme-enzyme interactions^10,11^. While compelling, these observations are typically restricted to specific pathways - such as glycolysis, the TCA cycle, secondary metabolites, or amino acid biosynthesis - and often involve multi-enzyme complexes with known physical interactions^7,11–13^. It was furthermore shown at the level of large-scale metabolic networks in relation to the associated network of protein-protein interactions (PPIs of enzymes), that paths in PPIs mirror metabolic network distances, suggesting an evolved structure-based co-optimization^5,14^.

Advances in protein structure prediction have opened new opportunities to move beyond binary protein-protein interaction networks in which proteins are treated as structureless interaction nodes, and to investigate the spatial organization of metabolism at the atomic level and at scale. Methods such as AlphaFold2^15^, AlphaFold3^16^, ESMFold^17^ can generate both reliable structural models of the involved proteins and accurate models of protein–protein complexes. Given the availability of reliable structural models generated by the three listed AI-modeling approaches, classical docking-based approaches, such as HDOCK^18^, can also be employed to explore potential physical associations among large sets of enzymes. Thus, the computational quantification of spatial relationships between functionally relevant features, such as catalytic sites, is now possible at unprecedented scale and resolution.

Here, we combine curated catalytic site annotations with metabolic pathway information to assess whether enzymes catalyzing sequential reactions in *Escherichia coli* tend to exhibit reduced spatial separation between their catalytic sites when modeled as interacting pairs, thereby reducing diffusion times and uptake probabilities of reaction products of one enzyme taken up by the next enzyme as substrates (Figure 1A). Using data from the Catalytic Site Atlas and KEGG pathway annotations, we identified 107 enzyme pairs involved in consecutive metabolic steps. For each pair, we generated structural interaction models using four independent prediction frameworks and quantified lower bounds on diffusion path lengths between catalytic sites using both Euclidean distances and physically more plausible SASP distances (Shortest Accessible Space Path distances).

**Figure 1.**
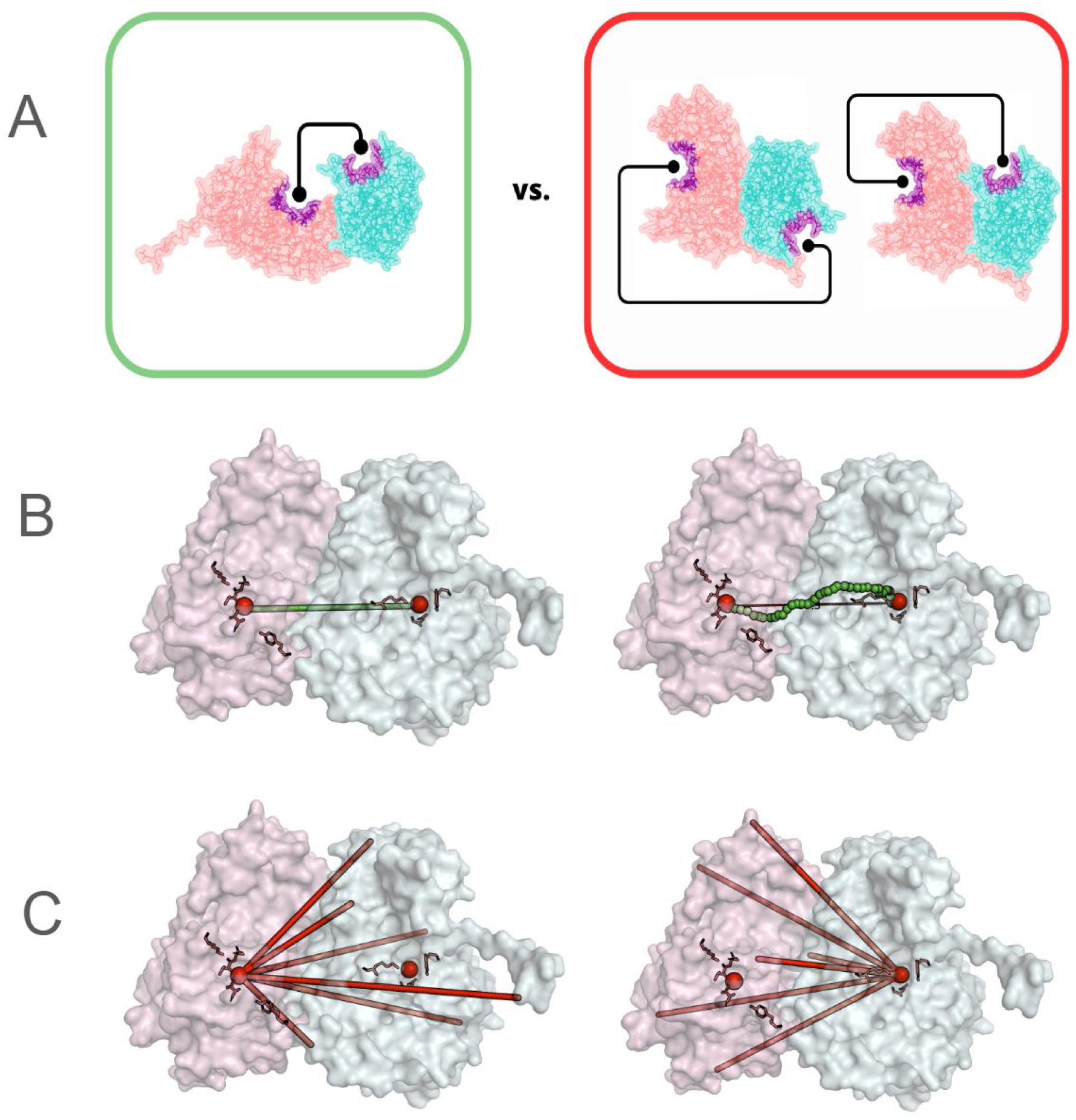
A) Hypothesis tested in this study: enzymes catalyzing subsequent metabolic reactions physically interact (often weakly), and in their docked conformation, they position their respective catalytic sites in closer proximity (left) than randomly expected (right). B) Illustration of Euclidean (left) and Shortest Accessible Space Path Distance (right) as the shortest, physically possible surface-traversing path between catalytic sites (green spheres). Red spheres represent the centroid of the annotated catalytic site residues (highlighted red)), of two interacting enzymes. C) Comparison of actual distances to catalytic site - random points on the respective other enzyme; likewise, this is done for Shortest Accessible Space Path distances.

To place these measurements in context, we compared observed catalytic site distances to appropriate random controls, including random surface points and random enzyme pairings. By systematically contrasting different modeling approaches and distance metrics, we aim to evaluate not only whether spatial proximity is observed, but also how robust such observations are to methodological choices.

Aside from well-documented cases of stable multienzyme complexes (metabolons) referred to above, it seems possible that physical enzyme-enzyme encounters may often be weak and transient. Thus, we first examine how well various structure and protein-protein interaction prediction methods capture such weak interactions.

Our results indicate that while predicted enzyme pairs frequently exhibit reduced Euclidean distances between catalytic sites relative to random expectations, this effect is substantially attenuated when diffusion paths are evaluated using physically more plausible accessible space path distances. Moreover, similar proximity statistics are observed for randomly paired enzymes when interaction is assumed. At least to some degree, this effect can be attributed to geometric effects associated with catalytic sites found as invaginated sites on protein surfaces.

Taken together, our findings do not support a universal optimization of metabolic enzyme organization for minimal three-dimensional diffusion distances, but they do point to localized instances of spatial proximity that may reflect pathway-specific adaptations. Our observations highlight both the potential and the limitations of structure-based approaches for studying the physical organization of metabolism and motivate further investigation into context-dependent enzyme organization.

## Results

### Benchmark set for low-affinity interactions

Transient interactions are by their nature short-lived, reversible, and dynamic, rendering them difficult to crystallize. Consequently, they are rare in the Protein Data Bank (PDB). To evaluate how well current computational models handle these weak interactions and, therefore, to assess their utility when applied to protein (enzyme) pairs of unknown structure whose interactions are often transient, a dataset of 112 low-affinity dimers was extracted from PDBbind v20202. These dimers were classified as weak interactions based on a binding-affinity threshold, with dissociation constants greater than 1 µM (pK_d_ < 6).

On this benchmark set, we evaluated the performance of four state-of-the-art prediction methods: AlphaFold2 (AF2), AlphaFold3 (AF3), ESMFold, and HDOCK, to correctly capture interaction structures and affinities, as reflected by the reported confidence scores.

The overall distribution of DockQ scores across the 112 low-affinity dimers (Figure 2) suggests variable success among the tested methodologies. DockQ is a continuous 0–1 score that evaluates how accurately a predicted protein–protein complex reproduces the native interface by combining iRMSD, lRMSD, and contact recovery into a single metric.

**Figure 2.**
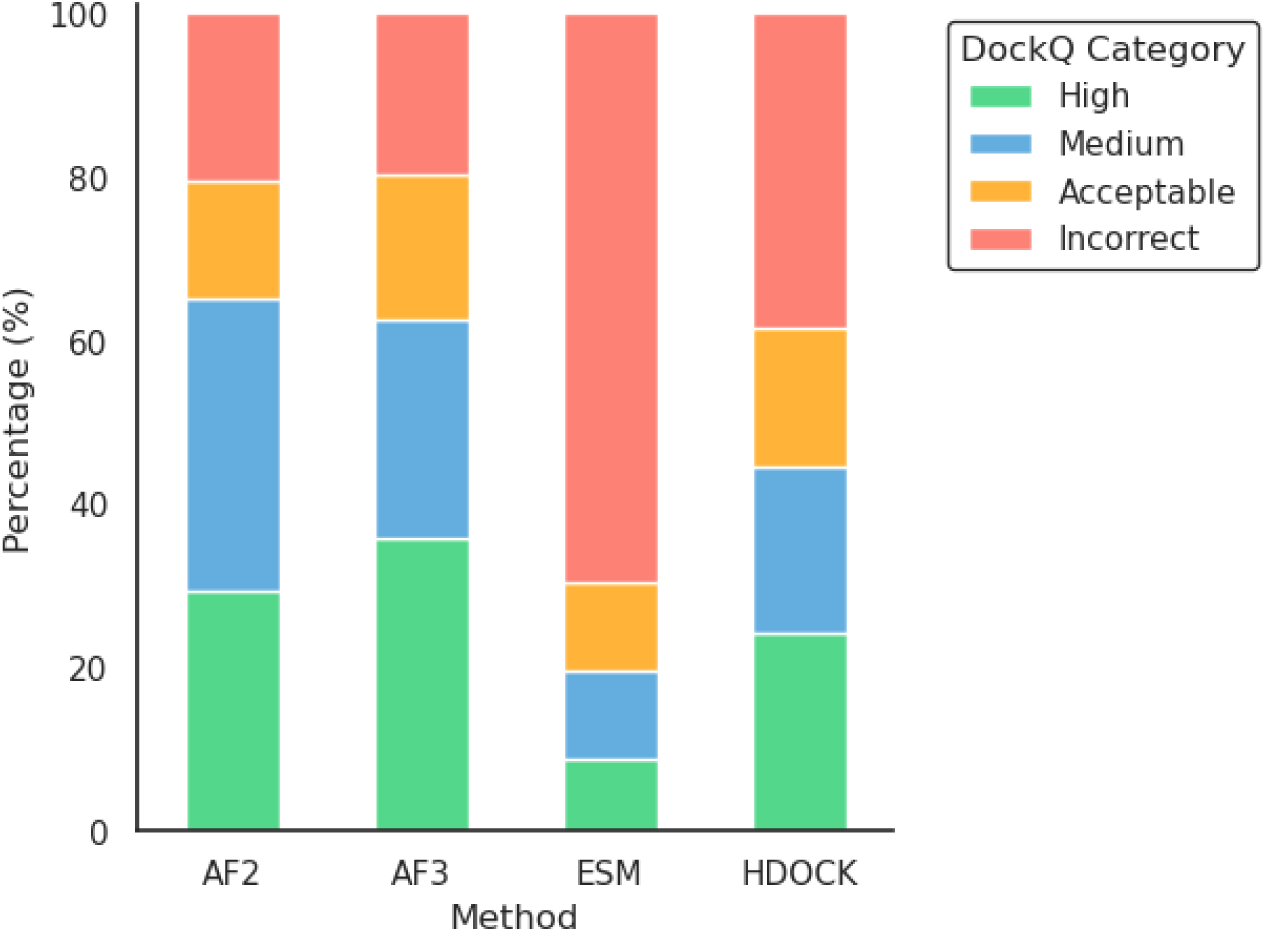
DockQ distribution of predicted complexes for four structure prediction methods on the benchmark set of 112 low-affinity protein dimers. The chart shows the percentage of models from AlphaFold2, AlphaFold3, ESMFold, and HDOCK that fall into four DockQ categories: Incorrect (DockQ < 0.23), Acceptable (0.23 to 0.49), Medium (0.49 to 0.8), and High (DockQ > 0.8).

AF2 and AF3 showed relatively good performance in predicting weak protein-protein interactions. Approximately 60% of the predictions fell into the medium or high quality categories, and 80 % into the acceptable medium or high categories. The classical rigid-body docking method, HDOCK, had a lower percentage of correct predictions, with approximately 60% predictions in the acceptable medium or high categories. ESMFold displayed a higher percentage of models categorized as incorrect (approximately 70%).

### Scoring methods for low-affinity interactions

Crucially for our study, we need to establish that the confidence scores reported by the various protein-protein interaction prediction methods can be reliably interpreted. While the available metrics have been well tested, since we are dealing with transient, low-affinity interactions, this relationship remains to be revisited. Hence, we investigated the correlation between DockQ and several single-model scoring functions on the benchmark set of weak interactions.

Different scoring procedures yielded different correlations, depending on the method used. The correlations of all available metrics for each prediction method are shown in Supplementary Figure 1.

The score ipTM yielded the highest correlation with DockQ values (Figure 3). This AlphaFold confidence metric, focused on the robustness of the interface region, showed a strong correlation with DockQ (r=0.82 for AF2 and r=0.74 for AF3). ipSAE also showed strong correlations with DockQ (r=0.78 for AF2 and r=0.69 for AF3) and it has been shown to perform better in protein structures with disordered regions^19^. However, these metrics are calculated using the AlphaFold PAE (Predicted Aligned Error) score. Thus, their use is restricted to AlphaFold predictions.

**Figure 3.**
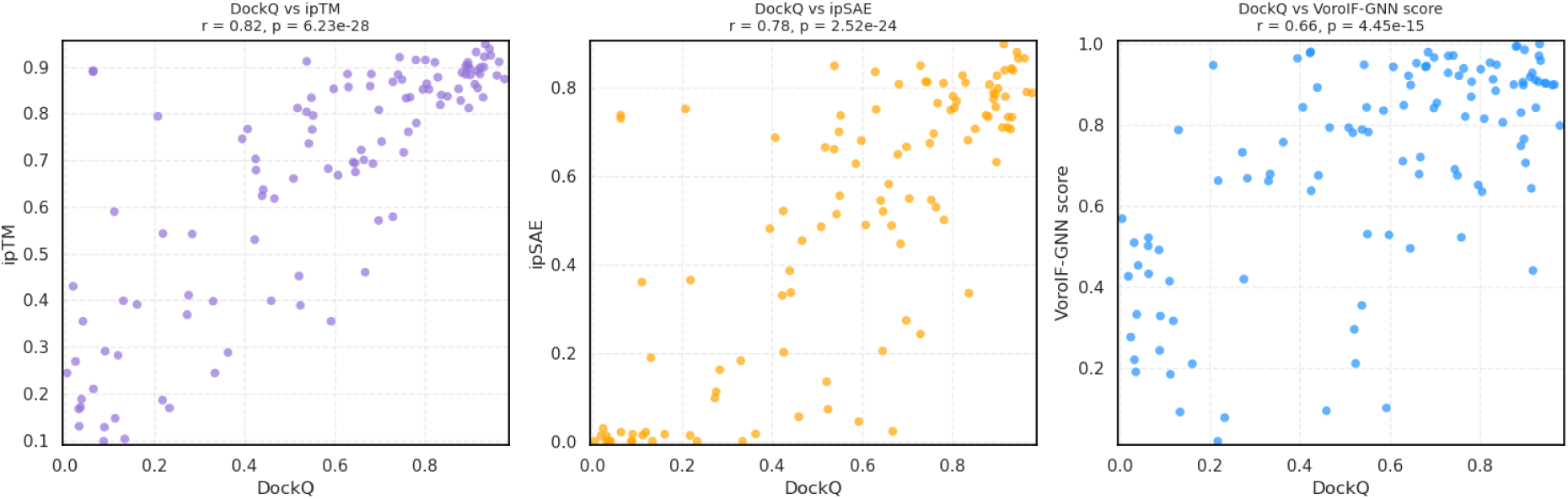
Correlations between the AF2-predicted protein dimers and associated ipTM, ipSAE, and VoroIF-GNN scores and their similarity to the reference structure (DockQ score).

We evaluated scoring functions independent of the AlphaFold Predicted Aligned Error (PAE) to enable quality assessment across diverse modeling platforms, including HDOCK and ESMfold. VoroIF-GNN yielded high correlation metrics (Pearson correlation coefficient, r > 0.6) for AF2, HDOCK, and ESMfold but only r=0.45 for AF3. The correlations of all available metrics for each prediction method are shown in Supplementary Figure 1.

The combined correlation obtained is r=0.69 (Figure 4). By contrast, VoroMQA results were poor (Supplementary Figure 1). Although VoroIF-jury is an ensemble ranking score (which depends on other predictions rather than representing an absolute quality score), it performed well in our approach for all models but ESMfold (see Supplementary Figure 1). VoroIF-GNN scores yielded medium correlations across the four modeling methods.

**Figure 4.**
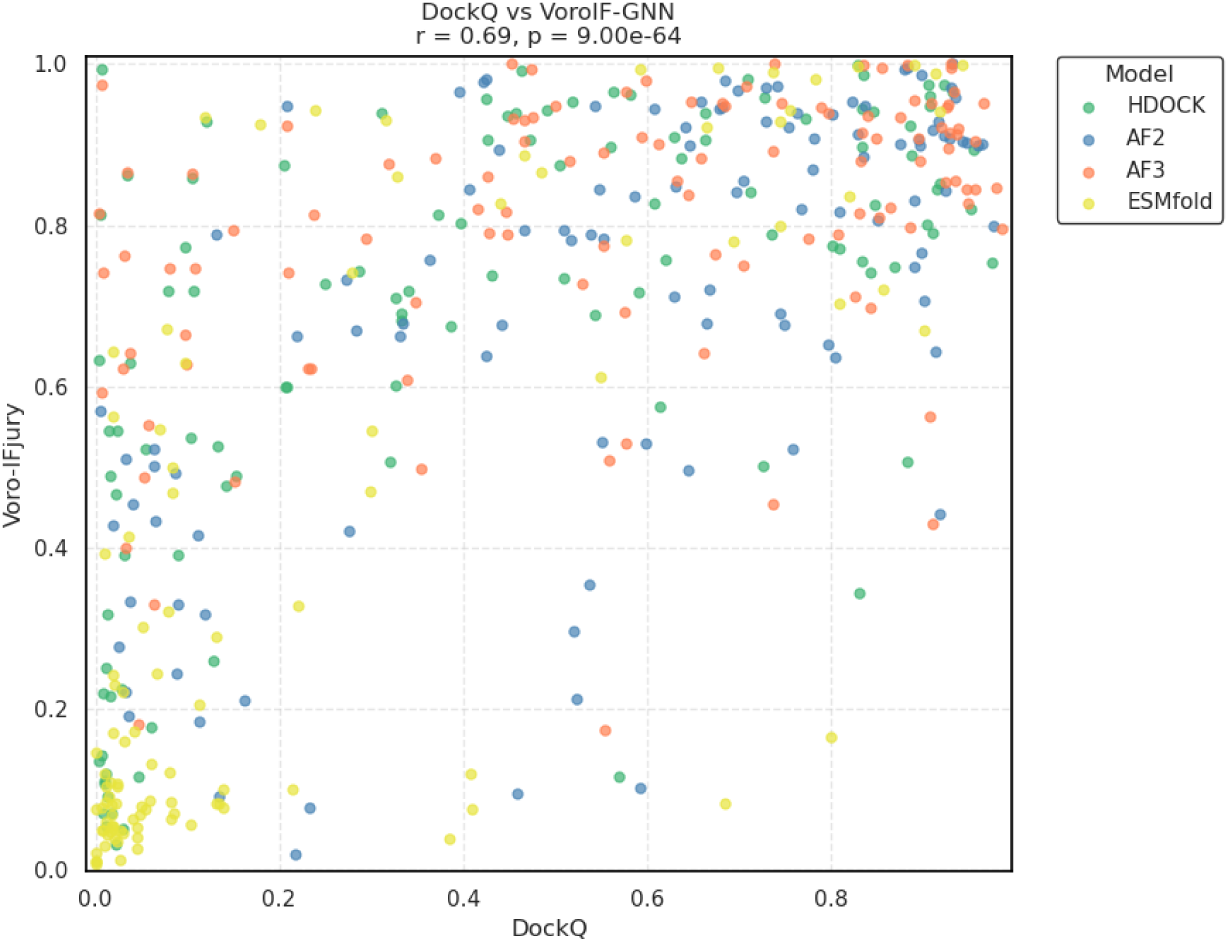
VoroIF-GNN correlation with DockQ similarity to the reference structure for each PPI prediction approach. HDOCK: 112 points, r = 0.64, p = 2.63e-14; AF2: 112 points, Pearson correlation coefficient, r = 0.65, p = 4.44e-15; AF3: 112 points, r = 0.45, p = 6.34e-07; ESMfold: 107 points, r = 0.75, p = 1.47e-20. The combined correlation obtained is 0.69, p=9.00e-64.

### Consecutive enzymatic step enzyme pairs

To investigate the physical organization of metabolic enzymes, 234 *Escherichia coli* proteins with annotated catalytic site information were extracted from the Catalytic Site Atlas. We chose *E. coli* because, for this organism, the available dataset was the largest and most comprehensive, followed by human (131 proteins), yeast (42 proteins), and others with even fewer annotated proteins https://www.ebi.ac.uk/thornton-srv/m-csa/stats/database/. By integrating this information with KEGG pathway annotations, we initially identified 269 enzyme pairs with the relation type “ECrel”, i.e., subsequent metabolic reactions involving enzymes with annotated catalytic sites. After removing redundancies arising from enzymes participating in multiple KEGG pathways, the final dataset comprised 107 unique enzyme pairs catalyzing sequential metabolic reactions.

These subsequent enzyme pairs map to different pathways (Supplementary Table 1). The most frequently represented pathways included “alanine, aspartate, and glutamate metabolism” (20 pairs) and “pyrimidine metabolism” (15 pairs).

All 107 pathway-sequential enzyme pairs were modeled as putative physical complexes using four independent structure prediction methods: AF2, AF3, ESMFold, and HDOCK. In line with the expectation that these pairs generally do not form stable complexes, the overall interaction confidence prediction scores were low (Figure 5).

**Figure 5.**
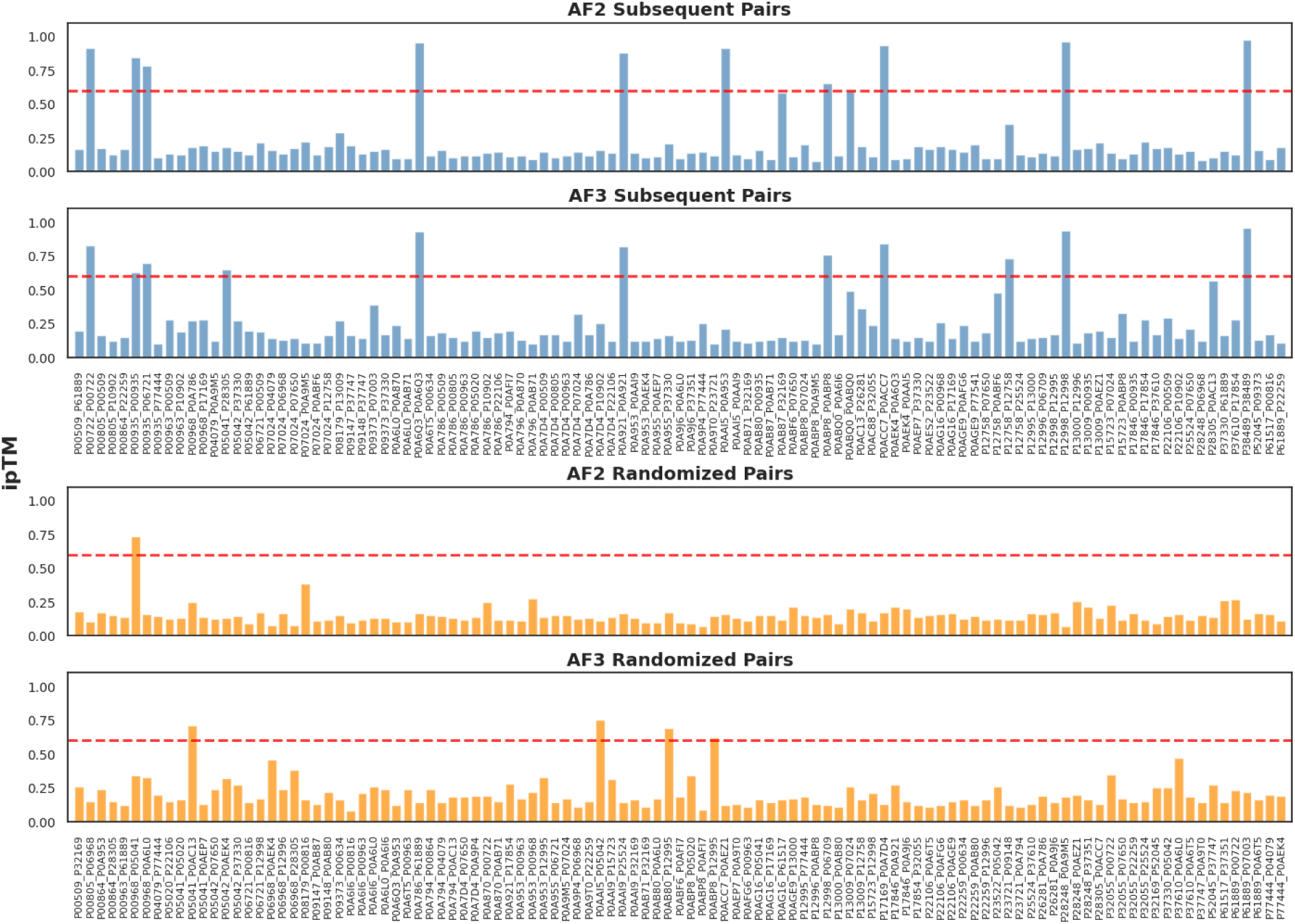
ipTM values for all predicted dimers predicted for subsequent enzymatic step enzyme pairs, and a same-size set of randomized pairings of those proteins using AF2 and AF3. The horizontal, dashed red line at ipTM=0.6 denotes the threshold above which predictions are considered high-confidence.

Across the four prediction frameworks, the variability in non-interaction outcomes reflects differences in how each method handles uncertainty. HDOCK, which relies on rigid-body docking, consistently produces interaction models because its search procedure enforces contact between monomers regardless of their true compatibility. By contrast, the cofolding approaches sometimes predicted no interaction between the enzymes. Among these, AF2 showed the most conservative behavior with 15 non-interacting enzyme pairs. It is followed by ESMfold, with 7 non-interacting protein pairs, and, lastly, AF3, with only one.

### Comparison of subsequent enzyme pair and randomized enzyme-enzyme interaction predictions

A direct comparison between methods is not straightforward, as each provides different interface confidence metrics. AF2 and AF3 report the interface predicted TM score (ipTM), HDOCK uses its own ITScorePP scoring function, and ESMFold does not provide explicit interaction confidence scores. Nevertheless, in AlphaFold predictions, when compared against randomly paired enzymes from the same dataset, subsequent enzyme pairs yielded higher ipTM scores (Figure 5). AF2 predicted 10 sequential pairs with ipTM > 0.6 (versus only one random pair), and AF3 predicted 11 sequential pairs with ipTM > 0.6 (versus four random pairs). From the high-confidence sequential pairs predicted with AF2, two present previously uncharacterized interactions (P00935_P06721 and P0AAI5_P0A953 (UniProt IDs)). Similarly, AF3 identified two uncharacterized interactions (P00935_P06721 and P05041_P28305).

Similar results were obtained with the internal scoring method of HDOCK, ITScorePP, where a baseline of low-confidence scores is shared between subsequent and randomized pairs, but in the subsequent enzyme pair set are more outliers (Figure 6). Applying a threshold of -350, 14 subsequent pairs obtained a more favorable score (more negative), in contrast to only four randomized pairs. From the high-confidence sequential pairs predicted with HDOCK, five present previously uncharacterized interactions (P00935_P06721, P00935_P77444, P0AAI5_P0A953, P0AC88_P32055, P25524_P07650).

**Figure 6.**
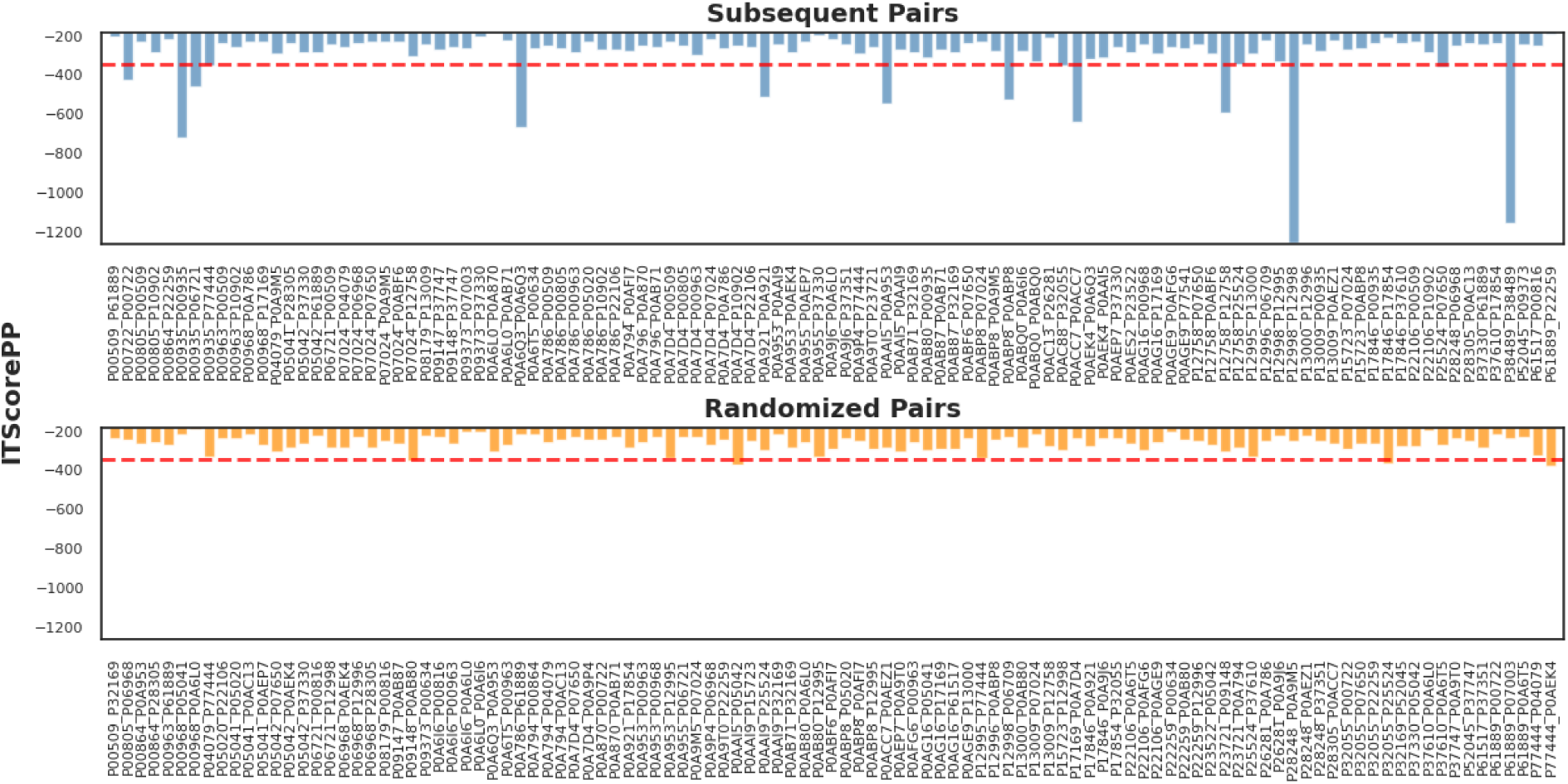
ITScorePP values for all predicted dimers of the subsequent pairs and randomized pairs sets using AF2 and AF3. The horizontal dashed red line at ITScorePP= -350 denotes the threshold below which predictions are considered high-confidence.

The score VoroIF-GNN was calculated for the four modeling approaches. Visually, the VoroIF-jury scores are consistently higher for sequential enzyme pairs than for random pairs (Figure 7), corroborating the trend observed with the AF2 and AF3 ipTM scores and the HDOCK ITScorePP score. Moreover, we observed that some enzyme pairs tend to score well across all the prediction methods. Interestingly, AF3 predictions yielded the highest VoroIF-GNN scores overall (with a mean of 0.3123).

**Figure 7.**
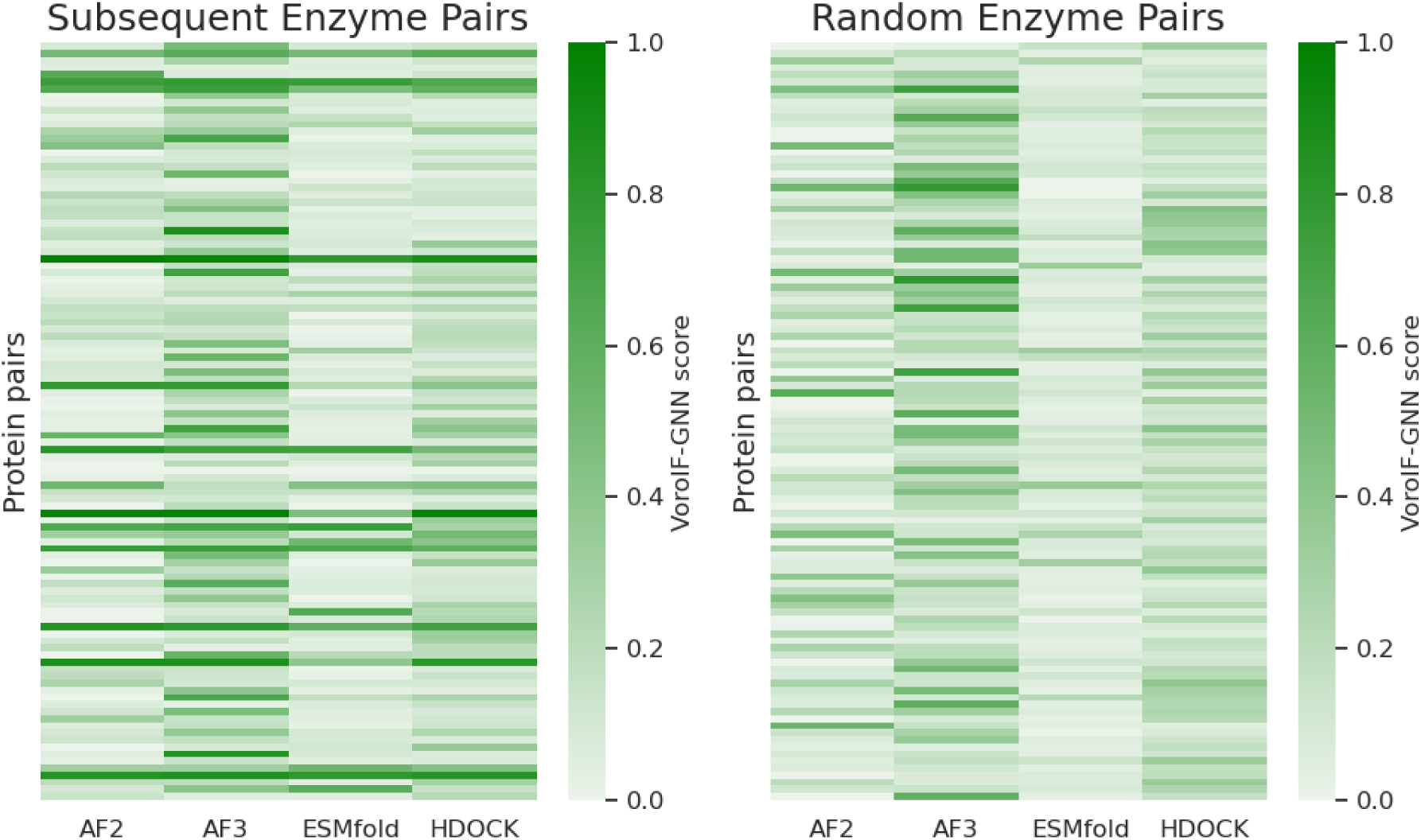
VoroIF-GNN scores for predicted enzyme–enzyme complexes. Heatmaps show jury scores (0–1) for (left) subsequent enzyme pairs and (right) randomly paired enzymes, across four modeling approaches (AF2, AF3, ESMFold, HDOCK).

Taken together, we established ipTM as a sensitive metric to gauge the validity of predicted conformations of weakly interacting proteins, an important step towards further interrogating the geometry of the mutual positions and orientations of catalytic sites on transiently/weakly interacting proteins. Furthermore, subsequent-step-catalyzing pairs are more often predicted to interact stably than random pairs within the same protein set (Figure 5).

### Euclidean distances between catalytic sites are consistently lower than randomly expected

For each modeled complex, we computed the Euclidean distance between the geometric centers (center of mass, same mass (one unit) assumed for all heavy atoms) of the annotated catalytic residues for the two enzymes. Distances were compared to those between the catalytic site of one enzyme and randomly sampled surface points on the partner enzyme.

Across all four modeling approaches, catalytic sites were predicted to be spatially closer (Euclidean distance) than expected at random. 78, 84, 91, and 85 out of 107 pairs supported this for AF2, AF3, HDOCK, and ESMFold, respectively (Figure 8).

**Figure 8.**
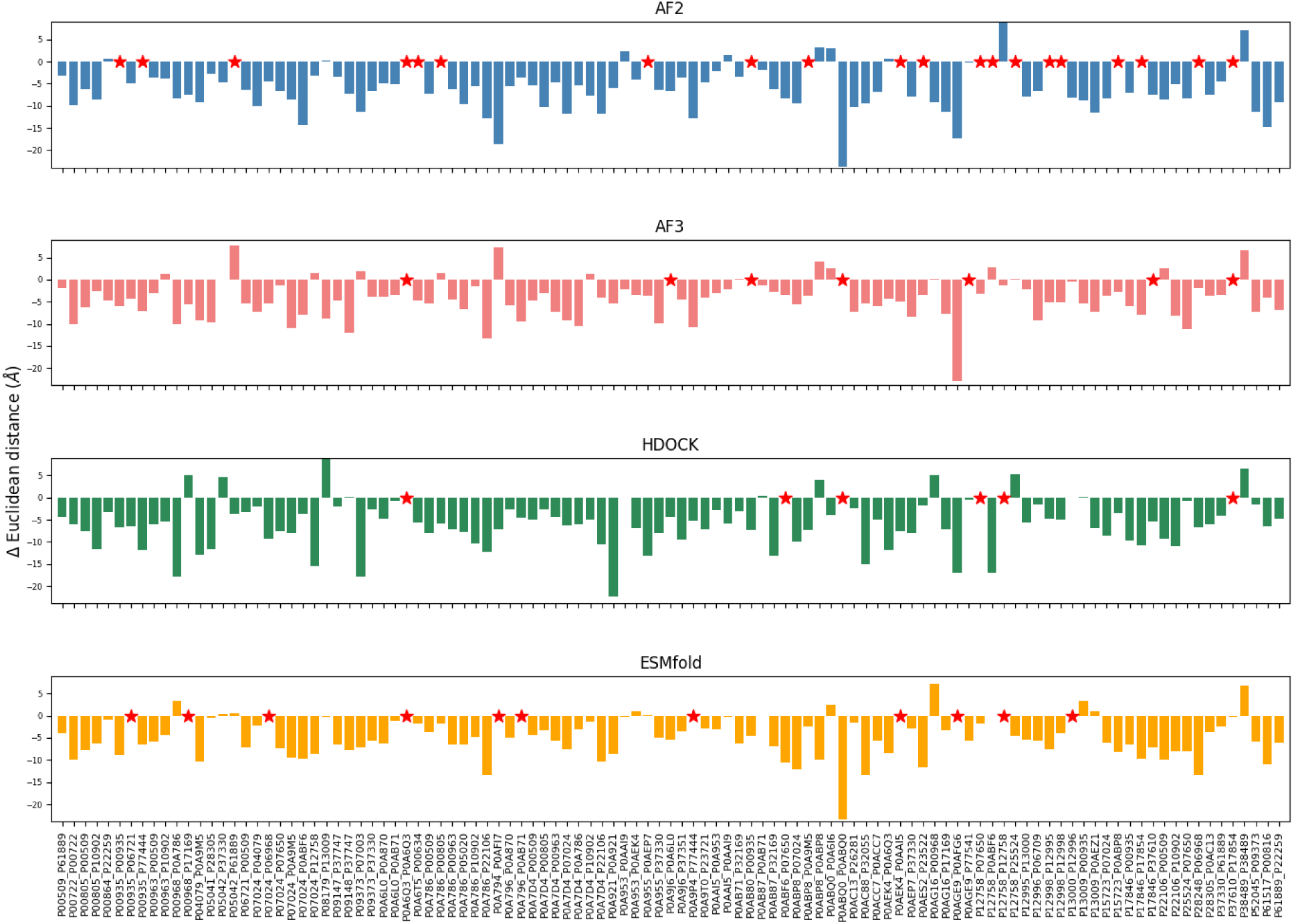
Difference, d_e_-d_r_, between the Euclidean distance of catalytic sites, d_e_, and the distance from catalytic sites to the random-surface-points median Euclidean distance, d_r_, for subsequent enzyme pairs. Evaluated using four different protein–protein interaction modeling approaches (AF2, AF3, ESMFold, and HDOCK). Red stars indicate that no interaction was predicted or that the catalytic site of one enzyme was blocked due to the interaction with the respective other protein.

When performing the same analysis on randomized enzyme pairings, a similar trend emerged. The majority of random pairs also showed catalytic site-catalytic site distances shorter than expected by chance. As shown in Figure 9, the distributions of catalytic distances for subsequent and random enzyme pairs substantially overlap across all four modeling approaches. The mean values for subsequent pairs were −6.56 Å (AF2), −6.20 Å (AF3), −5.12 Å (ESMFold), and −5.92 Å (HDOCK), while the corresponding random pair means were −6.20 Å, −3.63 Å, −5.12 Å, and −5.92 Å. While it is surprising that also in the case of random pairings, catalytic sites are closer to one another than randomly expected, to be discussed further below as a possible consequence of catalytic sites being found in locally depressed (invaginated) surface regions, except for HDOCK, all AI-based, co-folding methods for interaction prediction yielded larger negative difference values, suggesting that there is a weak signal for actual consecutive enzyme pairs to position their catalytic sites closer to one another than in the case of random pairing.

**Figure 9.**
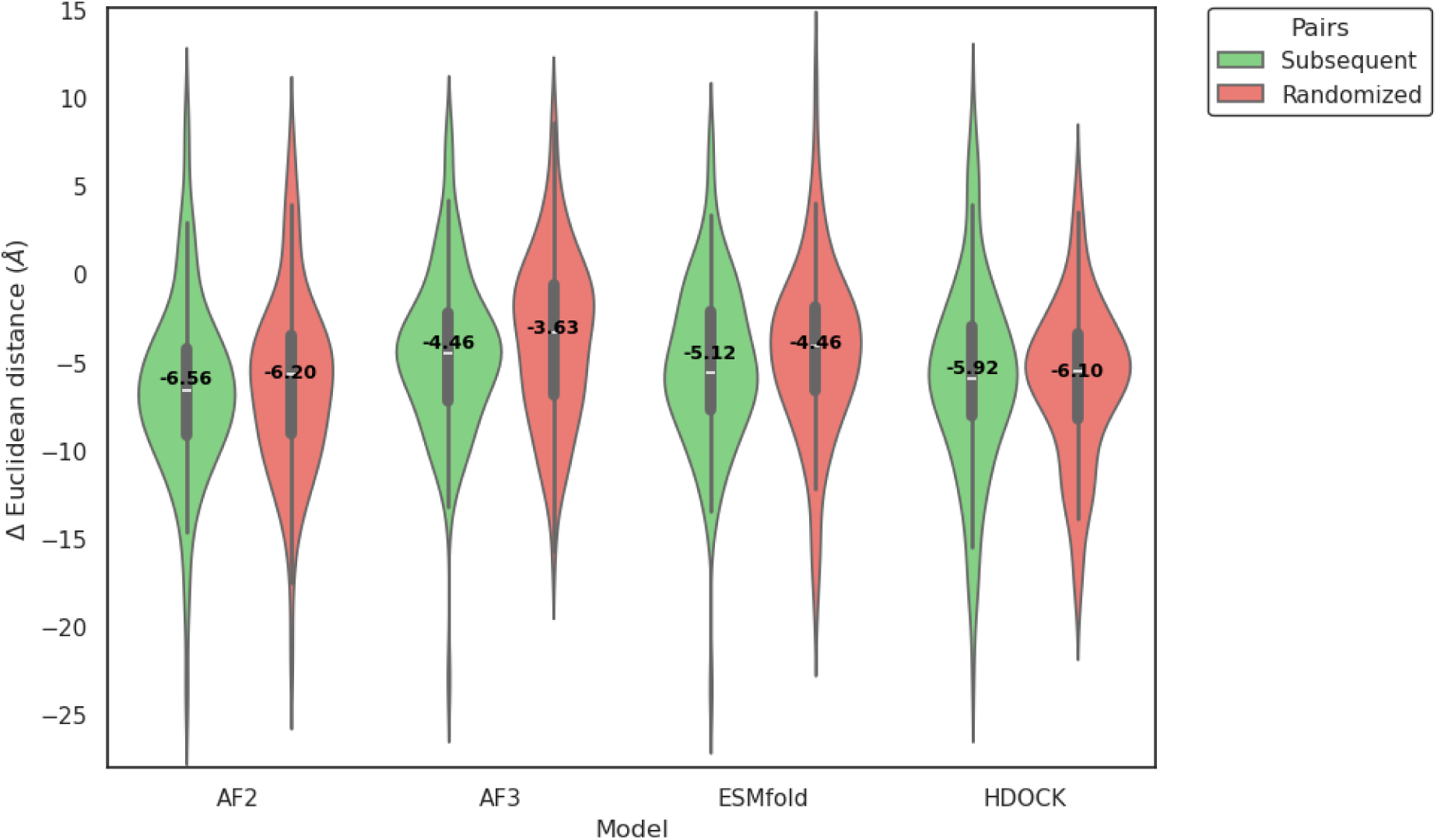
Comparison of Euclidean distance differences (catalytic site - catalytic site vs. catalytic site - random surface median) distributions between subsequent enzyme pairs and random enzyme pairs across the four modeling approaches.

We tested whether the site-distance statistic correlates with predicted interaction confidence scores, i.e. whether pairs with increased structural prediction confidence show a stronger signal of distance reduction. As for the confidence score, we used the VoroIF-GNN score, as i) it can be computed for all four protein interaction prediction methods used in this study, and ii) we observed that it meaningfully reflects the correctness of interaction structure prediction (Figure 3). However, when protein pairs were sorted by VoroIF-GNN score, no systematic trend was observed (no correlation) (Supplementary Figure 2). Thus, we cannot postulate that high-confidence interaction predictions show a stronger signal with regard to distance reduction (actual vs. random). Of note, we also inspected the ipTM-scores for AF2/3 predictions and arrived at the same conclusion.

### Shortest Accessible Space Path (SASP) distances: no evidence of spatial optimization

To realistically approximate the minimal physical path that a compound (metabolite substrate) would have to traverse along the protein surface and through the surrounding space, we calculated the shortest possible path, the Shortest Accessible Space Path (SASP), along the accessible space between catalytic sites in a dimeric conformation of two enzymes (see Methods), i.e. avoiding unphysical passes through atom-occupied space. The number of pairs with shorter SASP than expected by chance was lower than for Euclidean distance, with 53, 51, 56, and 70 out of 108 pairs for AF2, AF3, HDOCK, and ESMfold, respectively (Figure 10). Thus, no statistical signal of shorter-than-expected shortest SAS paths was detected.

**Figure 10.**
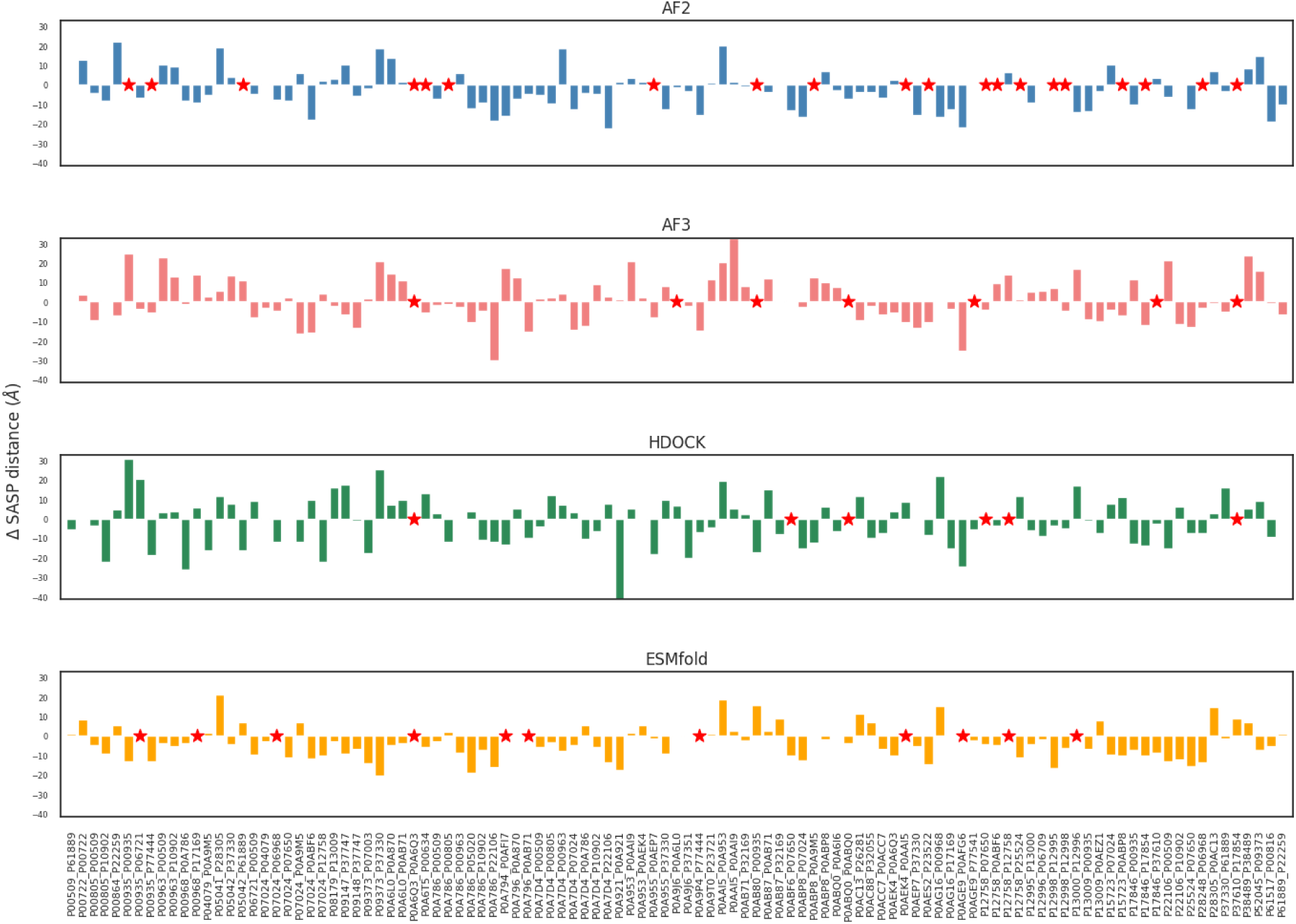
Difference, d_e_-d_r_, between the SASP (Shortest Accessible Space Path) distance of catalytic site pairs, d_e_, and the distance from catalytic sites to random surface points (median), d_r_, for subsequent enzyme pairs, evaluated using four different protein–protein interaction modeling approaches (AF2, AF3, ESMFold, and HDOCK). Red stars indicate that no interaction was predicted or that the catalytic site of one enzyme was blocked due to the interaction with the respective other protein.

As done for Euclidean distances, we tested whether the site-distance statistic correlates with predicted interaction confidence scores, i.e. whether pairs with increased structural prediction confidence show a stronger distance reduction. However, also for SASP distances, when protein pairs were sorted by VoroIF-GNN score, no systematic trend was observed (no correlation) (Supplementary Figure 3). As done before for the Euclidean distances, we also inspected the ipTM-scores for AF2/3 predictions and arrived at the same conclusion.

Randomized enzyme-pair interaction predictions showed that catalytic site distances were very similar to those of actual sequential pairs, despite lower interaction confidence scores. When comparing subsequent enzyme pairs to randomly paired enzymes, the distributions showed substantial overlap (Figure 11). Mean values for subsequent versus random pairs were very similar across models (e.g., −2.77 vs. −4.59 in AF2, 0.8 vs. −1.21 in AF3, -3.78 vs. −1.85 in ESMFold, and −1.45 vs. −2.96 in HDOCK). This indicates that, as with Euclidean distances, SASP-based differences do not clearly distinguish subsequent enzyme pairs from random controls, and no statistically significant separation between the two groups was observed.

**Figure 11.**
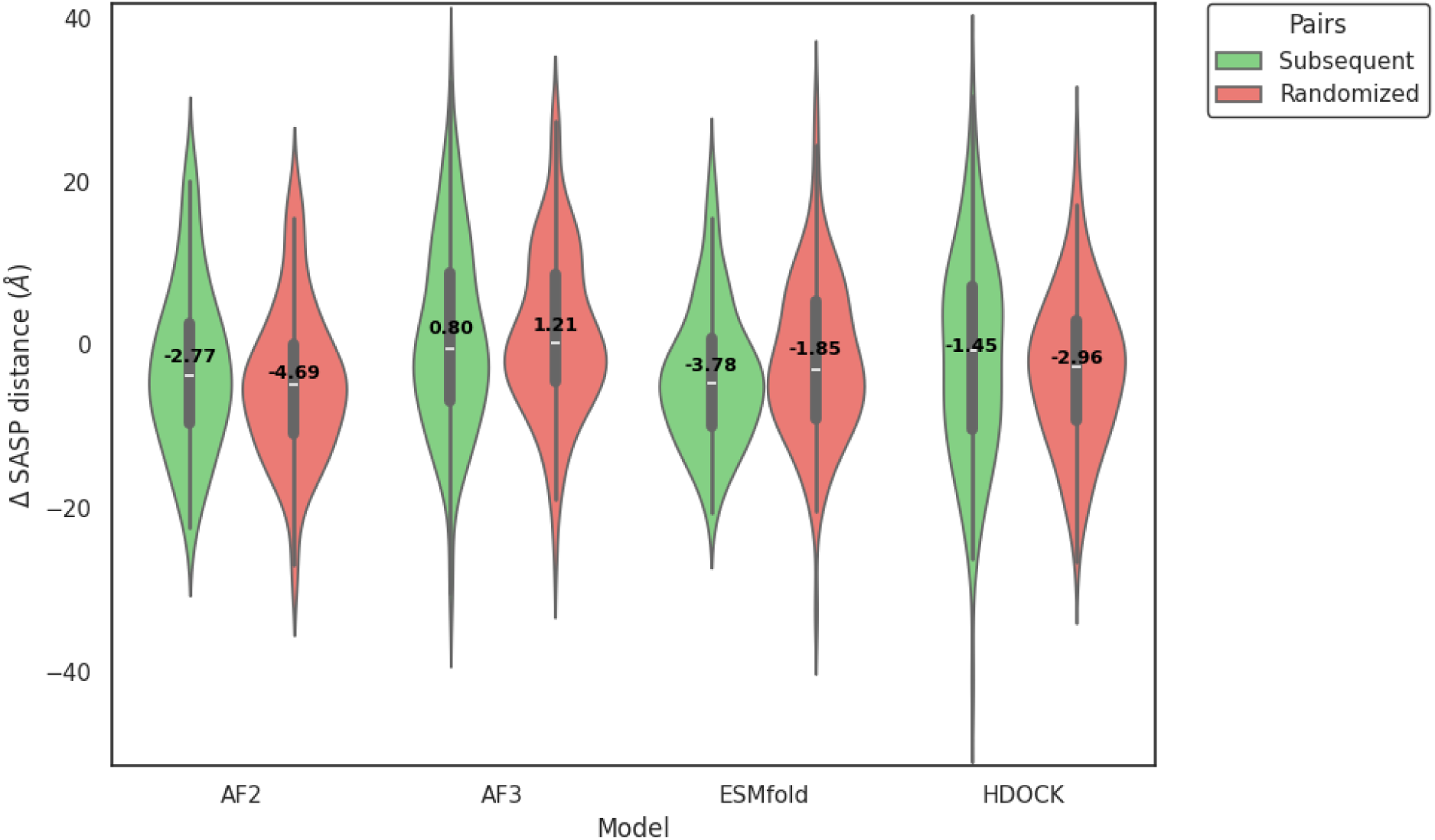
Comparison of SASP distance differences (catalytic site - catalytic site vs. catalytic site - random surface median) distributions between subsequent enzyme pairs and random enzyme pairs across the four modeling approaches.

### Effect of burial state of catalytic sites on distance statistics

The tendency for any protein-protein pair to position their catalytic site seemingly in closer proximity than randomly expected both when considering Euclidean (Figures 8, 9) and SASP (Figure 11, albeit less strongly), may originate from catalytic sites being generally found in invaginated surface regions, i.e. locally depressed surface regions (concave cavities as illustrated in Figure 1A). As illustrated and explained further in Supplementary Figure 4, assuming interacting proteins to be represented by ideal touching spheres of unit radius r=1, pairs of random surface points on the two spheres will be found, on average, at a Euclidean distance of 2.33. When assuming catalytic sites to be “buried” to different degrees, their mean pairwise distance will decrease, reaching 2, when located at the center of the spheres. It can be shown and was numerically verified (Supplementary Figure 4), that the average mean Euclidean distance, <ED>, goes with <ED>=2r+r/3*f^2^, where f is the burial factor (f=1: surface, f=0: center of the sphere/protein). Thus, assuming catalytic sites to correspond to burial factors f of 0.75, the <ED> will be 2.2, i.e. ∼6% less than that of the general protein surface positions (f=1). At f=0.5, the distance reduction is ∼10.7%. Assuming average proteins to have a radius of 20 Å, the relative reduction of distance computes as 2.4 Å and 4.3 Å, respectively, i.e. in the range of our observations (Figures 9). Similarly, SASP distances will be generally decreased for invaginated surface sites relative to general surface position as Euclidean and SASP distance can be assumed to be correlated to a certain degree (Figure 11), and much less consistently (Figure 10). Thus, we conclude that indeed the fact that catalytic sites are generally locally depressed surface regions contributes to their seemingly closer than random positioning.

### Novel, as of yet undescribed enzymatic complex

Most of the enzyme pairs that yielded high-confidence predictions are known to form stable complexes. However, one pair was found with no reported interaction (Figure 12). The enzyme pair corresponding to the UniProt identifiers P00935 and P06721 showed consistently high interaction scores across all modeling approaches (HDOCK ITScorePP = –460.97, AF2 ipTM = 0.79, AF3 ipTM = 0.70). As illustrated in Figure 12, the AF2 and AF3 models were highly concordant and positioned the catalytic sites in close spatial proximity, yielding a short SASP. By contrast, ESMFold and HDOCK generated markedly different complex arrangements. The HDOCK model did not support an optimized catalytic path, while the ESMFold prediction resulted in a blocked catalytic site, preventing construction of a feasible path between the two.

**Figure 12.**
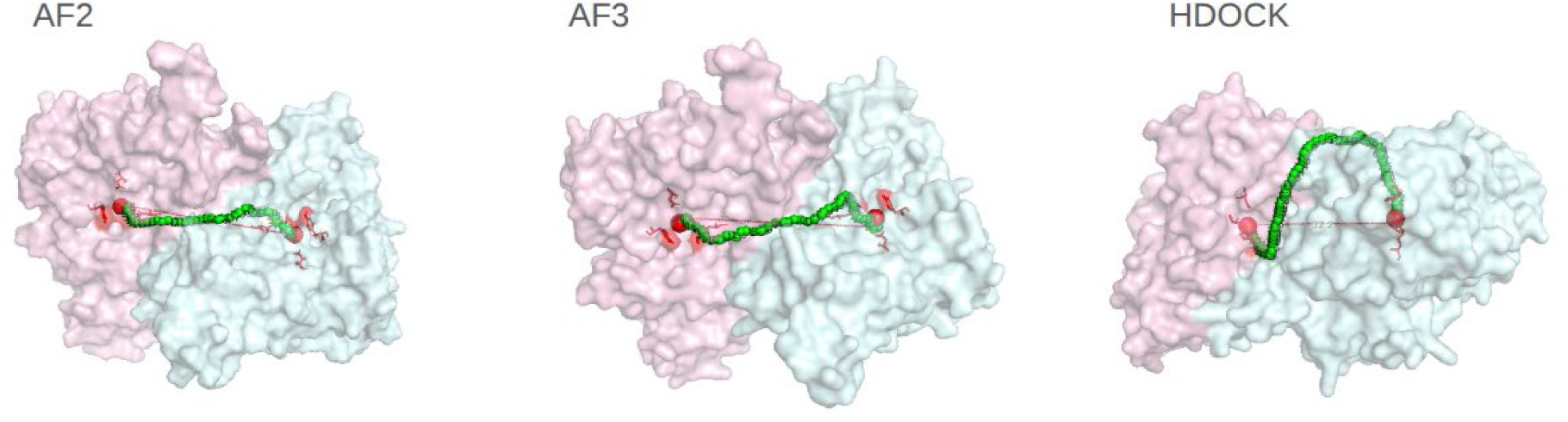
SASP path between the two catalytic sites of P00935-P06721 predicted interaction using AF2, AF3, and HDOCK. The pink protein corresponds to P00935, the blue to P06721, red residues are the annotated catalytic site residues, red spheres show the center of mass of each catalytic residue, and the trace of green spheres shows the SASP path.

These proteins correspond to MetB and MetC from *E. coli*, which function sequentially in the methionine biosynthesis pathway. MetB catalyzes the formation of L-cystathionine from O-succinyl-L-homoserine and L-cysteine, and MetC cleaves L-cystathionine into homocysteine, pyruvate, and ammonia.

Our results, together with favorable VoroMQA and VoroIF-GNN scores, support the potential existence of a previously uncharacterized interaction between these two sequential-step enzymes. The AF2 and AF3 models further suggest that their spatial arrangement may facilitate optimized substrate channeling between the catalytic sites.

## Discussion

In this study, we tested the hypothesis that if enzymes catalyzing consecutive metabolic reaction steps (transiently) interact, they position their respective catalytic sites in close proximity to increase throughput via shorter diffusion distances, leading to increased uptake probabilities by the “next” enzyme. We tested this hypothesis on 107 consecutive enzyme pairs in *E. coli* using state-of-the-art protein-protein interaction prediction methods. As a result, we found no evidence supporting our hypothesis. While Euclidean distances between catalytic sites were found to be shorter than between randomly chosen surface points on the same protein pair (Figure 8), the same was true for random enzyme-enzyme pairings (Figure 9), for which such “advantage” should not act as an evolutionary pressure, and possible directly attributable to catalytic sites being invaginated, concave surface pockets (see above and Supplementary Figure 4). More critically, when considering actual physically possible SASP distances, no difference between actual catalytic sites and random surface points was observed (Figures 10, 11).

Discussing the limitations of our study and their potential impact on the conclusions drawn, we see the following main aspects of our study that warrant consideration: 1) We selected *E. coli* as a representative species along with its metabolism. 2) We relied on computational methods to predict the interaction of enzymes and the conformation of protein complexes. 3) We tested simple distance metrics (Euclidean and Shortest Accessible Space Path distances), irrespective of the actual size of metabolites and their physico-chemical properties, as well as their interaction with the proteins along their spatial traversal. 4) Our hypothesis posits that enzymes catalyzing consecutive reaction steps physically, at least transiently, interact, which may not be generally the case. 5) Lastly, we also need to ask whether any reduction in diffusion distances would actually be relevant in the context of catalysis, i.e., which speed-up seems realistic and what impact it could have given typical catalytic rates of enzymes and would an evolutionary pressure on optimizing catalytic site positionings be plausible.

With regard to the choice of species, *E.coli* offers a well-annotated metabolism and, more importantly, the largest set of annotated catalytic sites (234 vs. 131 for the second-most extensively annotated species (human)). Thus, it is the best choice given data availability. We cannot think of any characteristic of *E. coli* (prokaryote, single cell) that would make it a species in which the tested hypothesis is less likely to be true. Selection for high growth rates will favor an optimized metabolism.

As for the computational protein-protein (enzyme-enzyme) interaction prediction, of the four chosen ones, three employ the recently established “break-through” AI technologies that allow for high-confidence structural prediction methods (AF2, AF3, ESMfold). Their key advantage in our context is that they predict folding and interactions in parallel, thus potentially accounting for interaction-induced structural changes, the ignoring of which has been considered one of the main drawbacks of conventional rigid-body docking methods. Even though AF2/3 are state-of-the-art and have been proven to reliably predict protein structures, correctly protein-protein interactions remain challenging^20^. In addition to the three AI methods, HDOCK was chosen as a robust method that represents the class of rigid-body docking approaches. Thus, we used the most current, best available computational methods.

As the central parameter for testing our hypothesis, we used the simple Euclidean distance between catalytic sites and the more involved Shortest Accessible Space Path distance. While simple, the Euclidean distance can only provide a crude estimate of actual path distances, as, evidently, straight lines between catalytic sites are almost never sterically possible. To address this limitation, we implemented and made available (https://github.com/joacolongo/SASP) a computational procedure to compute the shortest sterically possible path between two protein sites, the Shortest Accessible Space Path (SASP). While this accounts for the obvious limitation of Euclidean distances, as the SASP computation ignores any interactions between the metabolites and the proteins (local physical interactions with protein sidechains), and as the substrate was considered a probe of size solvent (water), ignoring the actual metabolites and their physico-chemical character, the spatially shortest SASP path may actually not be the best physical path. It has long been known and described for compound - protein-binding-site encounters^21,22^ that by “steered”-diffusion, i.e., “guided” by physical interactions, compounds are steered towards a target site. However, accurately accounting for such properties and interactions would render the study computationally prohibitive.

We operated under the assumption that enzymes catalyzing consecutive metabolic reaction steps, i.e., the product released by one enzyme is the substrate of the next, physically interact, at least transiently. Even though stable enzyme complexes that realize this assumption have been described (as outlined in the Introduction), this may not generally be the case, and enzymes may be freely diffusing in the cytoplasm. While we observed more protein pairs with high interaction scores for the true consecutive-step enzyme pairings than for random pairings (Figures 5, 6), our results on interaction scores (Figure 5) may be taken as evidence that, often, there are no appreciable effective interactions. There seems to be a binary classification of enzyme interactions: either stably interacting or not at all, or, if so, very weakly. Conspicuously, no intermediate score values were obtained. This may, however, reflect on the paucity of suitable training data. Protein-protein interactions have been trained exclusively on stable complexes, as for transiently interacting proteins, relevant structural information is not available to the same extent or at the same level of 3D resolution. At best, binary calls (interacting/not-interacting as obtained from cross-linking experiments and the like) are available^23^.

A critical consideration when interpreting the lack of shorter than random proximity between catalytic sites is whether a reduction in diffusion-path distances would confer any selective kinetic advantage. For most metabolic enzymes, the rate-limiting step is the chemical transformation (k_cat_) rather than the physical encounter between the enzyme and substrate^24^. When enzymes operate far from the diffusion limit, the time required for a substrate to diffuse between active sites is negligible compared to the time required for catalytic turnover^25^. Theoretical modeling of enzyme cascades suggests that while proximity can drastically enhance flux for “imperfect” enzymes with low efficiency, this benefit vanishes for enzymes with typical cellular parameters once a steady state is reached^26^. In such systems, the local substrate concentration remains equilibrated with the bulk phase, rendering the precise orientation of catalytic sites kinetically “invisible” to the overall metabolic flux^27^. This theoretical framework provides a kinetic rationale for recent observations by Bhattacharyya and co-workers, who examined protein-protein interactions (PPIs) within E. coli carbon metabolism^28^. Their study reported that transient protein-protein interaction interfaces - while capable of appreciably boosting metabolic flux - are frequently distant from the active sites of the participating enzymes^28^. While qualitatively in agreement with our findings, the latter conclusion presented in Bhattacharyya and co-workers was based on visual inspection of dihydrofolate reductase and was not further quantified and broadly tested. Here, we pursued a more rigorous quantitative approach to assess distances between catalytic sites in predicted protein complexes and compare them to properly chosen controls (random surface points). While Bhattacharyya et al. focused on broad within- and inter-pathway interactions (e.g., folate and purine biosynthesis), our more rigorous quantitative approach confirms this trend specifically for consecutive reaction steps. The fact that flux can be enhanced by “local crowding”, as reported by Bhattacharyya and co-workers based on reaction-diffusion simulations, even when active sites remain distant, and that as reported similarly high-flux carrying pathways form tightly knit protein interaction networks^5^, suggests that the primary advantage of such assemblies may not be the reduction of diffusion travel time. Instead, as these enzymes operate far from the diffusion limit, evolutionary pressure may still favor locally concentrating and sequestering substrates, or may relate to aspects of protein stability or regulatory sensitivity over the presumably negligible kinetic gains that would be provided by precise spatial channeling of catalytic sites^29^.

## Conclusions

While enzymes catalyzing consecutive biochemical reaction steps in *E. coli* were observed to more frequently physically interact than random enzyme pairs, within the limits of our study, we found no evidence of an optimized positioning of catalytic sites of enzymes when physically interacting. In the context of weakly and thus transiently interacting proteins and found ipTM, ipSAE, and VoroIF-GNN, to be best correlated with true conformations.

## Methods

### Low-binding-affinity dimers dataset

As of the time of this study, PDBbind v2020 database^30^ contained 798 protein–protein complexes with experimentally determined binding affinities. In order to gather weak protein interactions, we selected all entries with dissociation constants greater than 1 µM (corresponding to pK_d_ < 6) binding affinity. Among these, only dimeric complexes with gap-free structures were selected.

### Dataset of consecutive enzymatic steps

*Escherichia coli* proteins with information about the catalytic site residues were extracted from the Catalytic Site Atlas^31^. KEGG IDs for these enzymes were obtained using the conversion API endpoint^32^. Only KEGG IDs corresponding to the *E. coli* K–12 genome (organism code eco) were retained. Complete pathway descriptions in KEGG Markup Language (KGML) format were obtained for each enzyme, and enzyme-enzyme relationships were extracted by parsing the KGML. Relations of type “ECrel” were considered indicative of biochemical consecutiveness. Enzyme pairs were recorded only when both members belonged to the original input set (proteins with annotated catalytic sites). This set is referred to as the “Subsequent Pairs” set.

An additional protein-protein pair set, the “Randomized Pairs” set, was created, in which the individual enzymes from the Subsequent Pairs set were randomly paired as a control.

### Protein-protein interaction modeling

We employed four approaches to predict the complexes formed by subsequent and random enzyme pairs. Three methods are based on deep learning architectures that simultaneously fold and dock (AlphaFold2-Multimer, AlphaFold3, and ESMFold), and one is a rigid-body docking approach (HDOCK).

**AlphaFold2Multimer (AF2)**^33^ predictions were performed with version 2.3.2 using PDB templates, five different seeds, and three predictions per seed.

**AlphaFold3 (AF3)**^34^ predictions were performed using version 3.0.1 with five different seeds and PDB templates allowed.

**ESMFold**^35^ predictions were performed using default settings and a recycle number of 20. For AF2, AF3, and ESMFold residues located at the N- or C-terminal regions with pLDDT scores below 60 were excluded after the complex modeling.

For **HDOCK**^36^, input monomeric structures were obtained from the AlphaFold Database v6 (https://alphafold.ebi.ac.uk). Terminal residues below the same confidence threshold (pLDDT < 60) were removed before docking. Then HDOCK was run with default settings using the public executable files.

### Catalytic site distance calculations

Catalytic sites were defined according to the residue annotations as provided in the Catalytic Site Atlas. In cases where annotated residues were spatially distant, residues whose primary role was to bind or stabilize cofactors were excluded. For each catalytic site, the center of mass of all atoms belonging to the remaining catalytic residues was computed. To map each catalytic site onto the molecular surface, we extracted the solvent-excluded surface points of the dimer using MSMS^37^ and identified the nearest surface point to the center of mass of the catalytic site residues. Distances between catalytic sites were then quantified using both Euclidean distances and Shortest Accessible Space Path (SASP) distance measures, computed with our developed path calculation method (Figure 1B, 1C; detailed below).

For statistical comparison, we also generated a reference distribution by sampling 150 random solvent-accessible surface points from both enzymes in a modeled dimer. Distances from each catalytic site to the paired enzyme random points were computed using both Euclidean and SASP. The difference between the distance of the two respective catalytic sites and the median of the resulting 300 (catalytic site of protein 1 to 150 random points of protein 2, and the catalytic site of protein 2 to 150 random surface points of protein 1, respectively) randomized distances was calculated.

### Minimal diffusion path distance calculation - Shortest Accessible Space Path (SASP) distances

To quantify distances constrained to the exterior of the protein, i.e., to capture physically plausible minimal diffusion paths, we devised the Shortest Accessible Space Path (SASP) distance metric. SASP, implemented in Python, computes the shortest path between two points restricted to the solvent-accessible surface points and the exterior of the protein, thereby preventing traversal lines through the protein interior and approximating a geodesic distance along the protein surface and straight lines to bridge inter-protein free (no protein atoms) space.

SASP constructs a triangulated solvent-excluded molecular surface using MSMS with a resolution of 1 Å and a probe size of 1.4 Å. A grid of auxiliary points is sampled in the solvent-exposed region surrounding the protein surfaces. A graph is then built by connecting all pairs of points separated by less than 1.75 Å, including both surface-mesh vertices and external auxiliary points. Using this connectivity graph, the shortest path between two user-defined coordinates is computed using the A* search algorithm^38^, ensuring that the route follows the exterior surface rather than the molecular interior (Figure 1B, right panel). To improve path efficiency, a post-processing refinement step prunes redundant points. A point of the resulting path is discarded if a direct line can be drawn between its predecessor and a subsequent point without intersecting the molecular surface or entering the protein interior. This refinement shortens the path and eliminates unnecessary detours. The final SASP is calculated by summing up the Euclidean distances between connected points along the refined path. The algorithm also produces a PyMOL script to visualize the resulting path.

The developed code for SASP computations is available at: https://github.com/joacolongo/SASP.

Of note, similar calculations have already been applied to cross-linking studies: Solvent Accessible Surface Distance (SASD)^39,40^.

### Protein-protein interaction prediction evaluation

To assess the quality of predicted protein--protein interfaces, we employed multiple complementary methods that capture geometric, energetic, and solvent-accessibility features as well as similarity metrics when a reference structure is available:

● DockQ: An interface similarity score that compares predicted and reference complexes, combining fraction of native contacts, ligand root mean square deviation (RMSD), and interface RMSD into a continuous quality metric^41^.
● Interface VoroMQA pseudoenergy: single-model scoring method (only the atomic coordinates are required) used for assessing protein structures and protein complexes. It functions as a generic interatomic contact-area-based energy potential. The method utilizes Voronoi tessellation to assess protein structure quality based on interatomic contact areas^42^.
● VoroIF-GNN: A single-model method that constructs a graph from Voronoi-tessellation-derived residue-residue contacts and uses an attention-based graph neural network to predict per-contact accuracy. These predictions are aggregated into a global interface score for ranking complex models^43^.
● VoroIF-Jury: ensemble score for ranking in protein complexes modeling. Its overall scheme involves collecting models from all sources, scoring and ranking them using several interface-focused methods (including VoroIF-GNN and VoroMQA variants), and then calculating a maximum achieved CAD-score consensus value across supersets of top models for the final ranking^44^. It was calculated using the top-ranked models of each modeling approach.
● PyMOL steric score: A van der Waals interaction energy-based score that evaluates models according to their optimization of steric complementarity. Calculated using PyMOL.
● iSASA: The interface Solvent Accessible Surface Area, calculated as the buried solvent-accessible surface area upon complex formation using freeSASA^45^.
● ipTM: The interface predicted TM-score from AlphaFold, which evaluates the consistency of predicted inter-chain geometry. Higher values indicate greater confidence in the relative orientation and packing of the interacting subunits.
● ipSAE: The interaction prediction Score from Aligned Errors is a redesigned interface confidence score that corrects the known biases of AlphaFold’s ipTM by restricting the calculation to residue pairs with reliable inter-chain PAE values. This produces a domain-aware interaction score that remains stable in the presence of long disordered regions or non-interacting accessory domains^19^. It was calculated with a PAE cutoff and a distance cutoff of 10 Å.
● ITScorePP: distance-dependent, knowledge-based scoring function used to evaluate protein–protein docking poses^46^. It is the scoring function used by HDOCK.
● PRODIGY: A knowledge-based predictor of binding affinity that estimates ΔG and dissociation constant (K_d_) values based on interface composition and contacts^47^.
● PBEE: Protein Binding Energy Estimator is a machine learning ensemble model designed to predict the absolute binding free energies (ΔG binding) of protein–protein complexes directly from their PDB coordinates^48^.

## Data and software availability

The developed computational procedures for computing Shortest Accessible Space Paths (SASP) between two sites on proteins may be found useful and have been made available (https://github.com/joacolongo/SASP). AlphaFold2/3, ESMfold, and HDOCK predictions are available for the 107 *E. coli* consecutive and randomized enzyme pairs along with Pymol scripts that load the computed SASPs and Euclidean distance information between the annotated catalytic sites are available at https://doi.org/10.17617/3.TZQZ3B.

## Authors’ contributions

DW conceived the study. JA and DW planned the study design and general implementation. JA performed all computations, algorithm developments and implementations, data analysis and result statistic generation. JA and DW interpreted the results and wrote the manuscript.

## Ethics declarations

Not applicable.

