## Supplementary Material for "Is metabolism spatially optimized? Structural modeling of consecutive enzyme pairs reveals no evidence for spatial optimization of catalytic site proximity"

**Supplementary Figure 1.** Comprehensive visualization of the correlation of confidence metrics as computed by the various interaction prediction methods and DockQ scores, taken as the gold standard, obtained for the benchmark set of known, weak protein-protein interaction pairs.

AF2

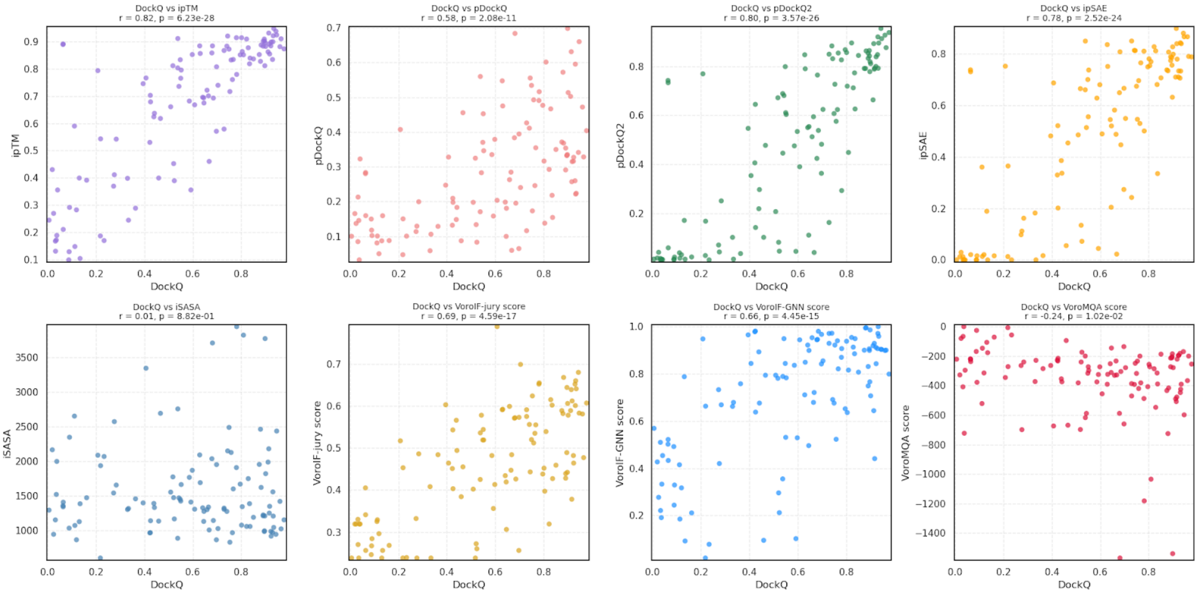


AF3

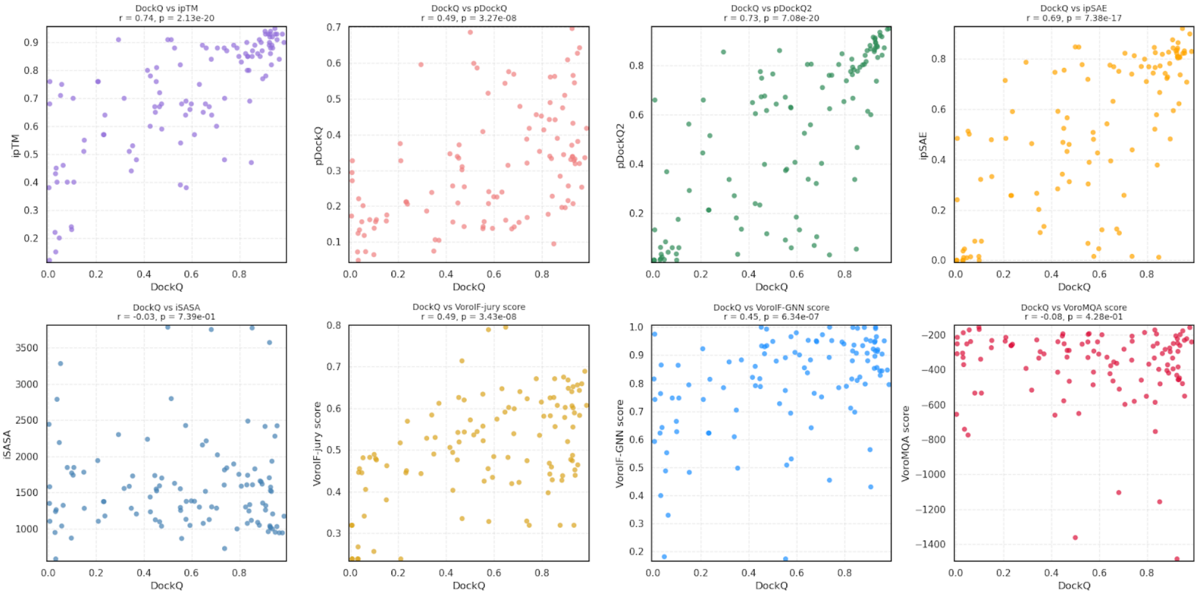


HDOCK

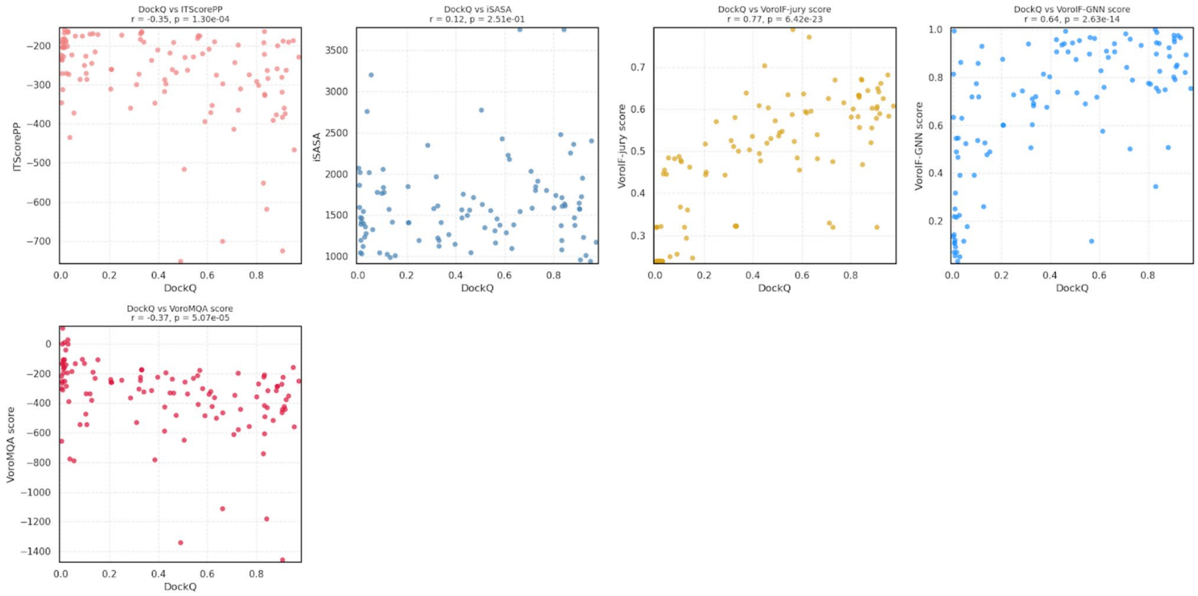
DockQ vs ITScorePP
r = -0.35, p = 1.30e-04
ITScorePP
DockQ
DockQ vs iSASA
r = 0.12, p = 2.51e-01
iSASA
DockQ
DockQ vs VoroIF-jury score
r = 0.77, p = 6.42e-23
VoroIF-jury score
DockQ
DockQ vs VoroIF-GNN score
r = 0.64, p = 2.63e-14
VoroIF-GNN score
DockQ
DockQ vs VoroMQA score
r = -0.37, p = 5.07e-05
VoroMQA score
DockQ

ESMfold

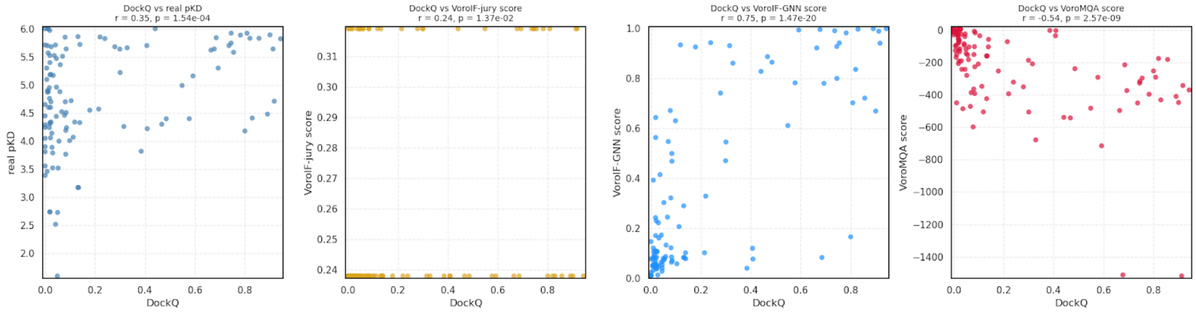
DockQ vs real pKD
r = 0.35, p = 1.54e-04
real pKD
DockQ
DockQ vs VoroIF-jury score
r = 0.24, p = 1.37e-02
VoroIF-jury score
DockQ
DockQ vs VoroIF-GNN score
r = 0.75, p = 1.47e-20
VoroIF-GNN score
DockQ
DockQ vs VoroMQA score
r = -0.54, p = 2.57e-09
VoroMQA score
DockQ

**Supplementary Figure 2.** Difference value, $d_e$-$d_r$, between the Euclidean distance of catalytic sites, $d_e$, and the distance from catalytic sites to the random-surface-points median Euclidean distance, $d_r$, for subsequent enzyme pairs. Evaluated using four different protein–protein interaction modeling approaches (AF2, AF3, ESMFold, and HDOCK), sorted by the VoroIF-GNN score in descending order, i.e. left-most pairs have highest score, i.e. higher confidence. Note that the sorting of pairs can be different for the four different interaction prediction methods. Red stars indicate that no interaction was predicted or that the catalytic site of one enzyme was blocked due to the interaction with the respective other protein.

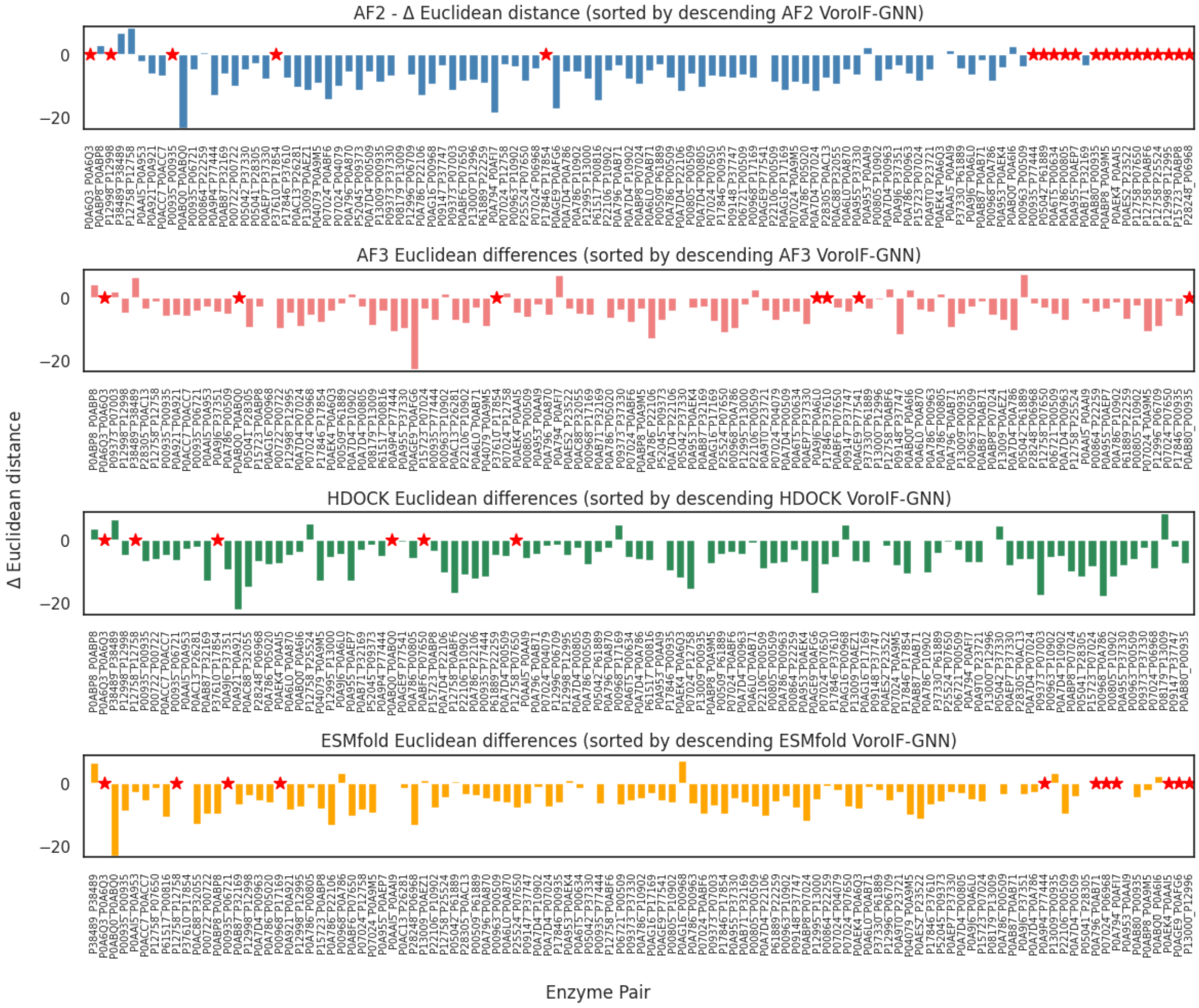

**Supplementary Figure 3**. Difference, $d_e$-$d_r$, between the ASD distance of catalytic site pairs, $d_e$, and the distance from catalytic sites to random surface points (median), $d_r$, for subsequent enzyme pairs, evaluated using four different protein–protein interaction modeling approaches (AF2, AF3, ESMFold, and HDOCK), sorted by the VoroIF-GNN score in descending order, i.e. left-most pairs have highest score, i.e. higher confidence. Note that the sorting of pairs can be different for the four different interaction prediction methods. Red stars indicate that no interaction was predicted or that the catalytic site of one enzyme was blocked due to the interaction with the respective other protein.

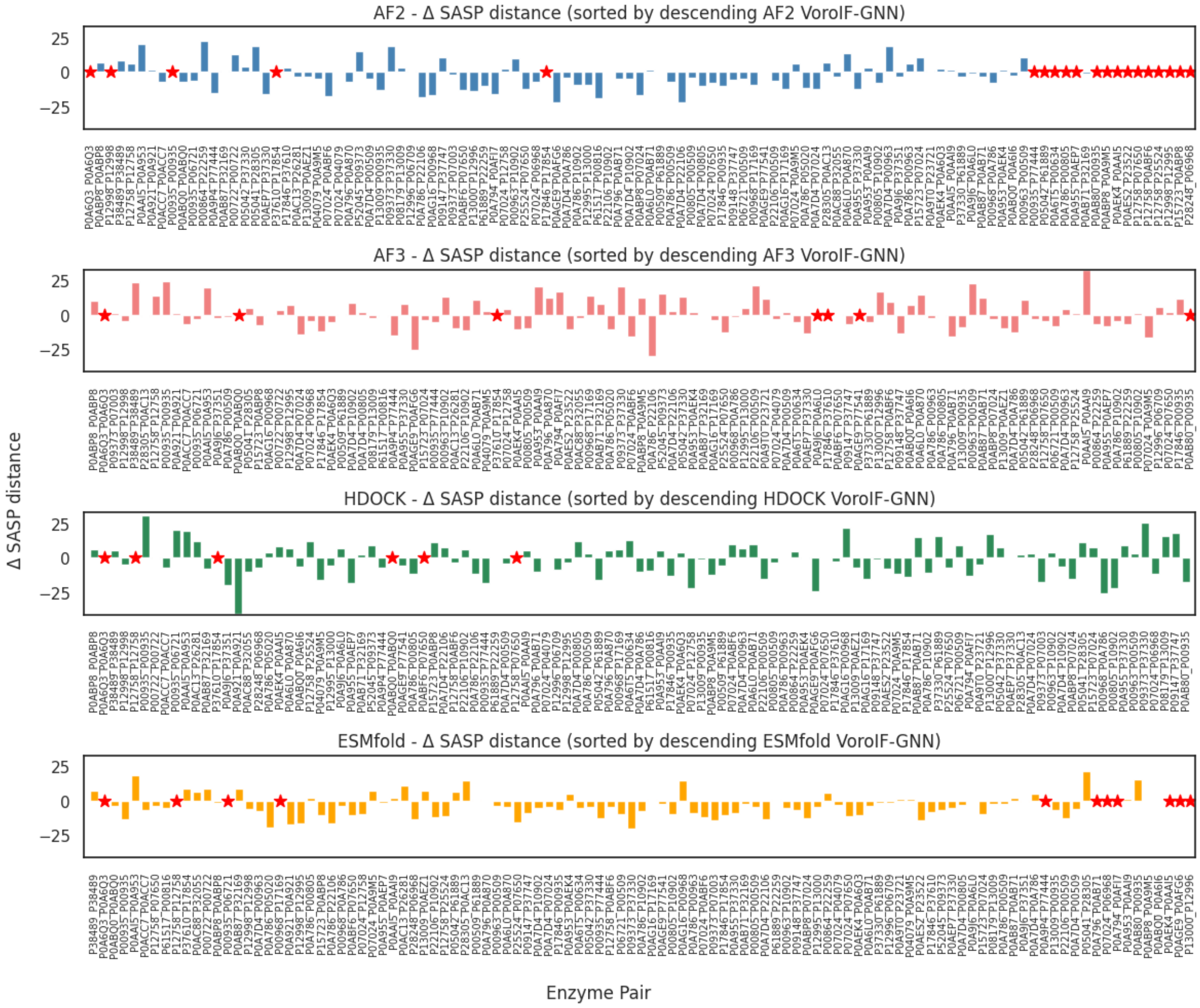

**Supplementary Figure 4.** Illustration of mean/median Euclidean distances of random surface points on two different touching spheres of equal radius (r=1, black dots), representing idealized and interacting (touching) proteins. The different concentric spheres represent different assumed burial depths of catalytic sites, with respectively chosen depth of burial, f*r, of f=1 (=surface of the sphere/protein, black dots), f=0.75 (red), f=0.5 (green) and f=0.25 (blue), respectively. The right-hand graph shows the numeric (simulated) values of means (circles) and medians (squares), compared to the theoretical values of <ED>=2r + r/3 * $f^2$ (grey, solid circles), where <ED> is the expected mean Euclidean distance between points on two same-color spheres. Numeric values were obtained from N=10,000 surface dots per sphere, uniform surface density, and 10,000 random sphere-sphere distances.

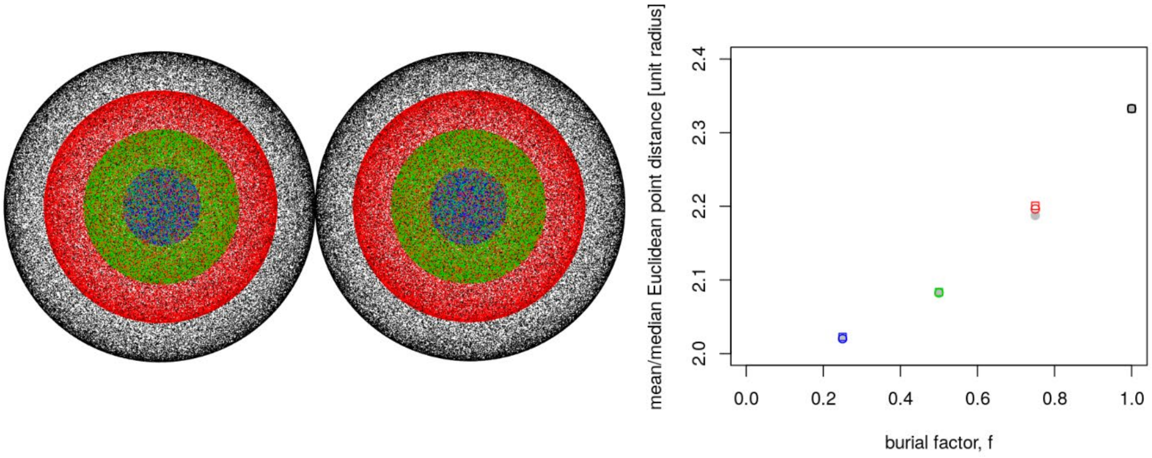

**Supplementary Table 1.** Coverage of KEGG pathways of the identified enzyme pairs catalyzing consecutive reaction steps.

| Pathway ID | Pathway_Name | no of pairs | UniProtID Pairs |
|---|---|---|---|
| eco00250 | Alanine, aspartate and glutamate metabolism | 20 | P00805_P00509 P00805_P10902 P00963_P00509 P00963_P10902 P00968_P17169 P0A786_P00509 P0A786_P00805 P0A786_P00963 P0A786_P10902 P0A786_P22106 P0A7D4_P00509 P0A7D4_P00805 P0A7D4_P00963 P0A7D4_P0A786 P0A7D4_P10902 P0A7D4_P22106 P0AG16_P00968 P0AG16_P17169 P22106_P00509 P22106_P10902 |
| eco00240 | Pyrimidine metabolism | 15 | P00968_P0A786 P07024_P06968 P07024_P07650 P07024_P0ABF6 P07024_P12758 P07650_P25524 P0A786_P05020 P0ABF6_P07650 P0ABF6_P12758 P12758_P07650 P12758_P0ABF6 P12758_P12758 P12758_P25524 P25524_P07650 P28248_P06968 |
| eco00061 | Fatty acid biosynthesis | 12 | P0A6Q3_P0A6Q3 P0A6Q3_P0AEK4 P0A953_P0AAI5 P0A953_P0AAI9 P0A953_P0AEK4 P0AAI5_P0A953 P0AAI5_P0AAI9 P0AAI5_P0AEK4 P0AAI9_P0A953 P0AAI9_P0AAI5 P0AEK4_P0A6Q3 P0AEK4_P0A953 |
| eco00230 | Purine metabolism | 12 | P04079_P07024 P04079_P0A9M5 P07024_P04079 P07024_P0A9M5 P07024_P0ABP8 P0A7D4_P07024 P0A9M5_P07024 P0ABP8_P07024 P0ABP8_P0A9M5 P0ABP8_P0ABP8 P15723_P07024 P15723_P0ABP8 |
| eco00270 | Cysteine and methionine metabolism | 9 | P00509_P61889 P00935_P00935 P00935_P06721 P06721_P00509 P06721_P00935 P0AB80_P00935 P13009_P00935 P13009_P06721 P23721_P0A9T0 |
| eco00780 | Biotin metabolism | 8 | P0AEK4_P0A953 P0AEK4_P0AAI5 P12995_P12998 P12995_P13000 P12996_P06709 P12998_P12995 P12998_P12998 P13000_P12996 |
| eco00030 | Pentose phosphate pathway | 7 | P0A6L0_P0A870 P0A6L0_P0AB71 P0A796_P0A870 P0A9J6_P0A6L0 P0A9J6_P37351 P0AB71_P0A796 P0AB71_P0A870 |
| eco00620 | Pyruvate metabolism | 7 | P00864_P61889 P05042_P37330 P05042_P61889 P09373_P07003 P09373_P37330 P22259_P61889 P61889_P37330 |
| eco00520 | Amino sugar and nucleotide sugar metabolism | 5 | P09147_P09148 P09147_P37747 P09148_P37747 P0ACC7_P0ACC7 P0ACC7_P27828 |
| eco00920 | Sulfur metabolism | 5 | P17846_P00935 P17846_P17854 P17846_P37610 P17854_P17846 P37610_P17854 |
| eco00051 | Fructose and mannose metabolism | 5 | P0A796_P0AB71 P0AB71_P32169 P0AB87_P0AB71 P0AB87_P32169 P32055_P0AC88 |
| eco00790 | Folate biosynthesis | 4 | P05041_P28305 P0A6T5_P00634 P0AC13_P26281 P28305_P0AC13 |
| eco00450 | Selenocompound metabolism | 4 | P00935_P77444 P06721_P00935 P0A9P4_P77444 P13009_P06721 |
| eco00052 | Galactose metabolism | 4 | G0ZKW2_G0ZKW2 P09147_P09148 P37747_P09147 P37747_P09148 |
| eco00710 | Carbon fixation by Calvin cycle | 4 | P00864_P22259 P00864_P61889 P0AB71_P0A870 P22259_P61889 |
| eco00630 | Glyoxylate and dicarboxylate metabolism | 4 | P0A955_P0AEP7 P0A955_P37330 P0AEP7_P37330 P37330_P61889 |
| eco00670 | One carbon pool by folate | 3 | P08179_P13009 P0AEZ1_P13009 P13009_P0AEZ1 |
| eco00020 | Citrate cycle (TCA | 3 | P0AGE9_P0AFG6 P61889_P05042 P61889_P22259 |

| | | | |
|---|---|---|---|
| | cycle) | | |
| eco00680 | Methane metabolism | 3 | P0AB71_P0A796 P23721_P0A9T0 P61889_P00864 |
| eco00770 | Pantothenate and CoA biosynthesis | 2 | P0ABQ0_P0A6I6 P0ABQ0_P0ABQ0 |
| eco00720 | Other carbon fixation pathways | 2 | P61889_P00864 P61889_P05042 |
| eco00541 | Biosynthesis of various nucleotide sugars | 2 | P0AC88_P32055 P0ACC7_P27828 |
| eco00640 | Propanoate metabolism | 2 | P0AGE9_P77541 P52045_P09373 |
| eco00300 | Lysine biosynthesis | 2 | P0A6L2_P04036 P0A9Q9_P0A6L2 |
| eco00261 | Monobactam biosynthesis | 2 | P0A6L2_P04036 P0A9Q9_P0A6L2 |
| eco00633 | Nitrotoluene degradation | 1 | P38489_P38489 |
| eco00564 | Glycerophospholipid metabolism | 1 | P0A921_P0A921 |
| eco00750 | Vitamin B6 metabolism | 1 | P0A794_P0AFI7 |
| eco00760 | Nicotinate and nicotinamide metabolism | 1 | P07024_P0ABP8 |
| eco00260 | Glycine, serine and threonine metabolism | 1 | P0A9T0_P23721 |
| eco00053 | Ascorbate and aldarate metabolism | 1 | P0AES2_P23522 |
| eco00910 | Nitrogen metabolism | 1 | P61517_P00816 |
| eco00010 | Glycolysis / Gluconeogenesis | 1 | P0AB71_P0A796 |

57
